# Basolateral Amygdala Represents and Remembers Ethological Events

**DOI:** 10.1101/2021.08.23.457318

**Authors:** Cristina Mazuski, John O’Keefe

## Abstract

The basolateral amygdala plays a crucial role in memory consolidation yet the general neural mechanism remains elusive. Basolateral amygdala neurons were recorded from freely-moving rats as they interacted with different ethological stimuli: male or female rats, a moving toy and food. Over 20% of neurons showed highly tuned *event-specific* responses to a single class of stimuli. Firing persisted in 30% of these responsive cells for minutes after the removal of the eliciting stimulus. Neural information flowed directionally from event-specific neurons to less specific neurons with changes in connection strength after removal of the stimulus. We propose that the basolateral amygdala identifies specific ethological events, with circuit-wide activity driven by the event-specific neurons during and after the termination of those events likely facilitating active short-term memory consolidation.

---

The basolateral amygdala has recently been assigned many different roles ranging from fear conditioning(*1*) and valence association(*2*, *3*) to exploratory behavior(*4*) and internal states(*5*). At the single-cell level, BLA neurons have largely been studied in the context of trained task-related activity, displaying responses to various cues including neutral sensory information (e.g. auditory, tactile, olfactory)(*6*), fear and pain cues(*7*, *8*), food(*9*) and conditioned valence responses(*3*). The presence of these small responses in broad, overlapping populations of BLA neurons has led to the suggestion that the complex, multi-sensory characteristics of naturalistic stimuli are likely to be encoded at the BLA circuit level as an aggregation of these low-amplitude noisy single-cell responses(*10*–*12*). In contrast, an earlier view proposed that single units could selectively identify events of significance to the animal(*13*). To distinguish between these alternative hypotheses, we recorded from large populations of neurons across the BLA using Neuropixels probes during several salient ethological events.

We report that subsets of BLA neurons consistently and selectively responded with large increases in firing activity in response to specific salient ethological events. Each of these distinct ethological events (presentation of male conspecifics, female conspecifics, a remote-controlled toy, and food) activated a distinct cluster of anatomically segregated neurons. Furthermore, approximately 1/3 of the activated cells showed persistent elevated activity long after the removal of the stimulus which continued to identify that stimulus. Detailed video analysis showed that the micro-behavioral aspects of the encounter such as which parts of the animals’ bodies touched further modulated the elevated firing activity. At the sub-millisecond level, many BLA neurons exhibited strongly cross-correlated activity, with information flowing from neurons tuned to one stimulus to those that responded to many stimuli and the strength of this connectivity increasing after the removal of the stimulus. These results show that single BLA neurons strongly represent distinct salient ethological events, convey this information to other BLA neurons and, in some cases, maintain a trace of these events long after their occurrence which might support active short-term memory for the event. In contrast, specific micro-behaviors result in weaker modulation at the single-cell level and are more strongly represented at the BLA population level.

We recorded from 426 neurons throughout the basolateral complex (Fig. 1A) of 5 male rats using single-shank Neuropixels probes. Each session consisted of a series of events (Fig. 1B and S1, Movie S1-4) during which the implanted rat could freely interact with an ethological stimulus placed in the recording chamber. Electrophysiological and video recording was continuous during a series of 5-min interactions with several males, females, a toy or sweetened rice food with 5-min baseline recordings before and after each event. We classified different cell types on the basis of their waveforms, sensory/behavioral correlates, and location within the basolateral complex, and will discuss the involvement of each of these cell groups in different aspects of the events.

**Fig. 1.**
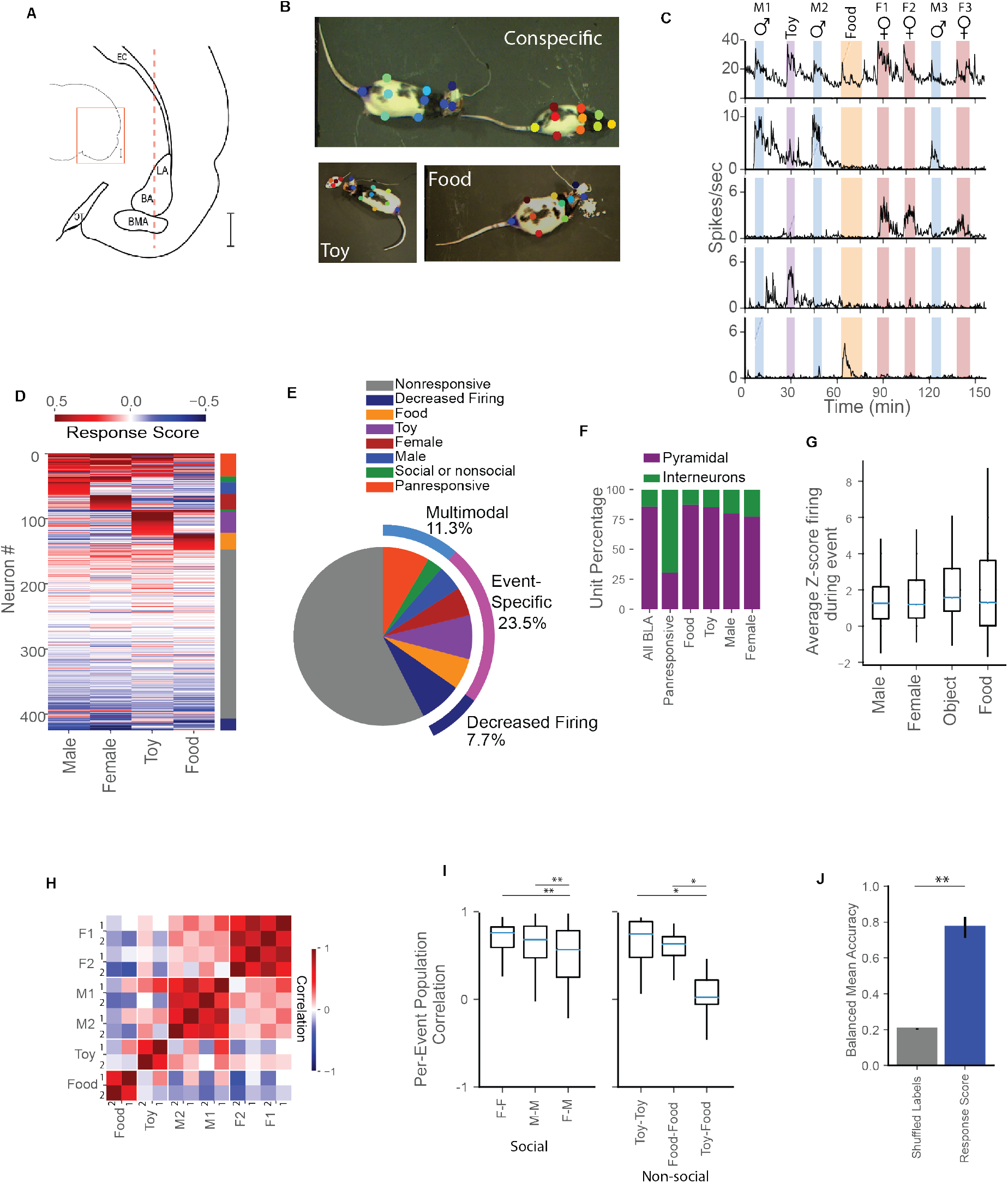
BLA Neurons respond to salient social and non-social stimuli. (**A**) Schematic of a coronal rat brain section depicting a Neuropixel probe (red dashed line) implanted in the BLA. (B) 10 body parts per rat were tracked from the recorded videos of social interaction using a combination of color-marking and Deeplabcut. Following post-processing, features from both the implanted rat and conspecific were extracted from these coordinates (see Movie S1-S4 and methods for more detail). Upper, Social interaction with a stimulus rat; lower left, interaction with a mobile toy mouse; lower right, eating sweet rice. (C) Each recording session consisted of 5 min Event-blocks of free interaction with one of 4 classes of stimuli interspersed with 5 min control periods. Stimuli consisted of (top) up to 3 different males (♂), up to 3 females (♀), a toy mouse (toy), and sweetened rice (food). Below, BLA single-unit activity reveals diverse responses in subsets of neurons. Representative categories of responses seen in the BLA: top to bottom – panresponsive, male-specific, female-specific, toy-specific, and food-specific. (D) Average response score for all BLA single units ordered by response strength and category. top to bottom – panresponsive, social: male-specific, female-specific, non-social: toy-specific, food-specific, no responses and decreased firing. (E) Percentage of BLA neurons with consistent responses to different events. (F) Panresponsive neurons are more likely to be putative interneurons than other event types. (G) There were no differences in the average z-scored firing activity of responsive neurons across different event types. (p = 0.32 One-way ANOVA. N = 306, 231, 134, 109 trials for male, female, toy and food respectively). (H) Population correlation of response score vectors from all responsive units across individual stimulus presentations shows strong correlations within specific events and weaker correlations across events. Each row represents first (1) and second (2) presentation of a given event-type (I) Quantification of population correlation for each event-type. (* p< 0.05, ** p< 0.01, N = intra-rat correlation of paired events, Kruskal-Wallis test with Dunn’s correction, N = 79, 30, 112 for social event pairs M, F, M-F, respectively, n= 4, 4, 16 for non-social event pairs Toy, Food, Toy-Food respectively). (J) Stimulus identity can be reliably decoded from neuronal response scores using LDA analysis. (** p<0.01, Student’s paired t-test, N = 4 rats)

## Results

### Single BLA cells identify the ethological event

The most striking result was that 23.5% of the cells showed a strong specific excitatory response to only one of the four classes of ethological stimuli (see Fig. S2 and methods for details). Fig. 1C shows typical examples of these *event-specific cells* and quantifies the proportion of cells responding to each event (Fig. 1D,1E). Typically, event-specific cells had a very low resting firing rate (median, 0.5 spikes s ^-1^) which increased many fold (3-6x on average) during the triggering ethological event. Importantly, the event-specific cells were highly selective responding to only one class of event, for example to all three females or to both males but showing virtually no response to other events (Fig. 1C,1D and S2). We found no evidence that the cells could discriminate amongst the different females or amongst the different males but cannot rule out the possibility that they might do so with further experience.

In addition to the event-specific responses, another 11.3% of the cells were multimodal, responding to more than one ethological event. Most of these neurons were *panresponsive*, responding to both social and non-social events (64% of panresponsive neurons responded to 3 or more events), and the majority of these were classified as putative interneurons based on their narrow waveforms and high firing rates as contrasted with the wider more silent putative pyramidal neurons (Fig. 1F, S3). A smaller proportion of neurons that were not classified as panresponsive or event-specific instead decreased in firing to one or more stimuli (7.7%). There were no significant differences in response intensity to different events suggesting that as a whole the BLA does not prefer one type of ethological stimulus over another (Fig. 1G).

At the population level, we observed strong population correlations between presentations of the same event-type, despite significant behavioral variability during the event-period (Fig. 1H, 1I). The selectivity of these responses for stimulus class were so strong that it was possible to use a decoder to identify the stimulus with 77% accuracy (Fig. 1J). Decoding of stimulus identity was possible with relatively few neurons (Fig. S4, performance plateaus with 10 neurons) and that effect was largely, but not exclusively, driven by the contribution of event-specific neurons (Fig. S4).

### Different stimuli are represented in different subdivisions of the BLA

The large number of electrodes and their density along the Neuropixels probes, together with careful placement of the probes within the BLA, allowed simultaneous recording from 2 or more subdivisions of the BLA including LA, BA and BMA in each animal (Fig. S5 and methods). All three divisions contained roughly the same percentage of multimodal panresponsive cells (11-12%) and the same percentage of cells that decreased firing (7-8%), but differed in other ways especially with respect to the other cell types. LA contained fewer responsive neurons, fewer putative interneurons and had lower average baseline firing rates compared to the BA and BMA (Fig. 2A, S5). Event-specific non-social (food and object) events were more highly represented in the LA and BA (77 & 67% of event-specific neurons) while conspecific events were more prominent in the BMA (74%) (Fig. 2B, C, S5). At the population level, this resulted in BMA and to a slightly lesser extent BA distinguishing strongly between males and females (Fig. 2D-E). Decoding of social stimuli improved at greater depths corresponding to the ventral BA and BMA (Fig. 2F).

**Fig. 2.**
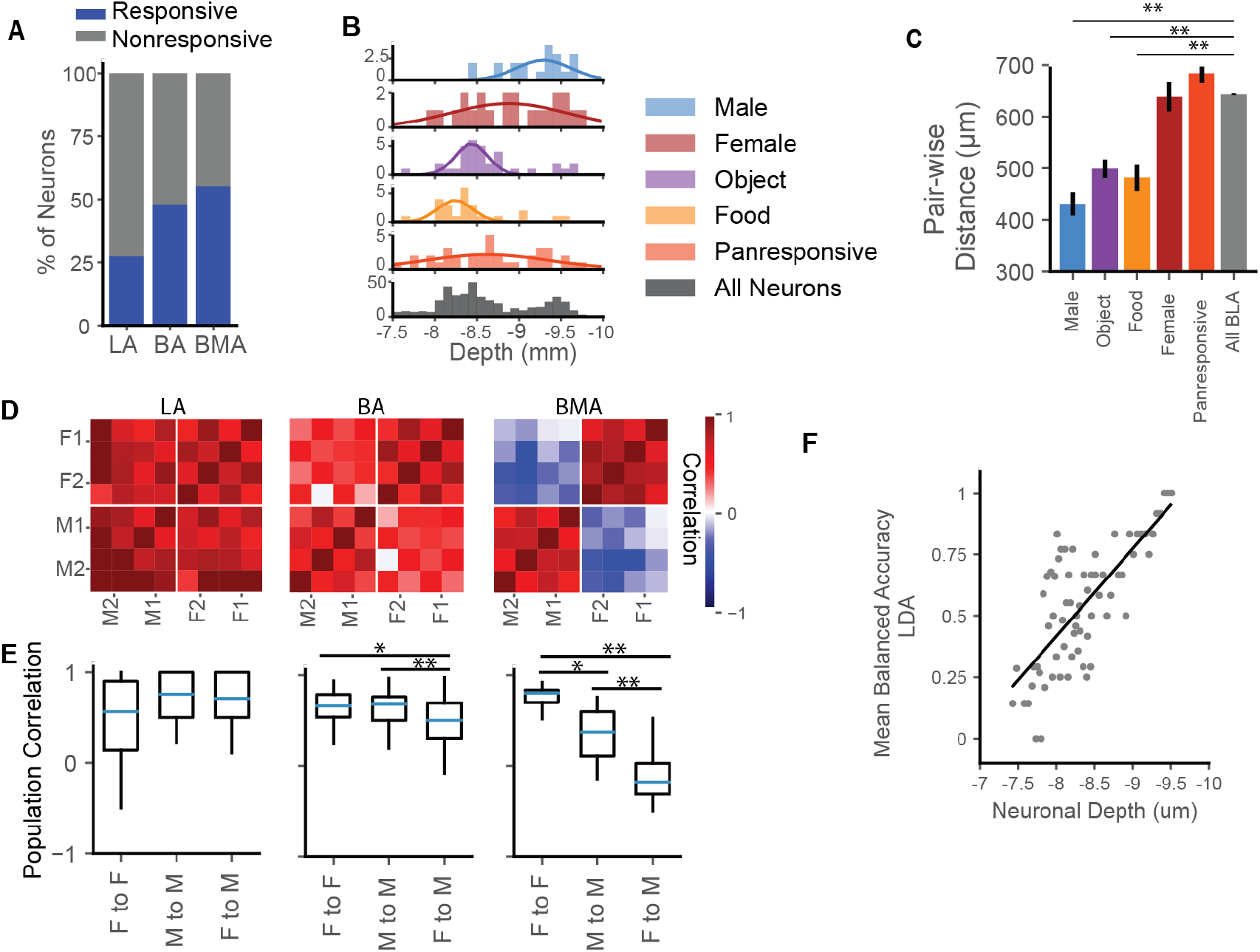
Regional Differences in BLA Stimulus Tuning. (**A**) A greater proportion of neurons in BA and BMA exhibit consistent responses to events than in LA (**p < 0.01, p = 0.1208, p = 0.4220 for percentage of LA, BA and BMA responsive neurons, compared to whole population, two-tailed binomial test). (B) Most event-specific neurons, specifically male, object and food but not female-responsive neurons, are clustered along the dorsal-ventral axis with (C) lower pair-wise distances than the average neuron. In contrast, panresponsive neurons are not clustered. (**p<0.01, pair-wise distance between pairs of neurons, N = 20, 34, 24, 22, 36, 426 neurons for male, object, food, female, panresponsive, and all BLA neurons, Kruskal-Wallis test with Dunn’s correction) (D) Population response vector correlation shows little discrimination between male and female conspecifics in LA, and greater discrimination in BA and BMA. This effect is quantified in (E): LA, p = 0.62 one-way ANOVA, BA *p < 0.05, **p < 0.01 one-way ANOVA, BMA *p < 0.05, **p < 0.01 Kruskal-Wallis test with Dunn’s correction, N(LA) =36, 12, 32, N(BA) = 24, 58, 84, N(BMA) = 15, 15, 36 for event-pairs F-F, M-M and F-M, respectively. (F) Decoding of social stimulus identity changes along the dorsal-ventral axis, with better discrimination of social conspecifics at deeper co-ordinates. (accuracy of decoding social events, R^2^=0.56, **p < 0.01, simple linear regression).

### Increased firing in responsive neurons temporally linked to identification of the ethological event

Responsive BLA units exhibited large sudden increases in firing activity around the time of event onset (Fig. 3A, 3B). To further understand the temporal dynamics of this increase, we divided the event start into distinct periods (presentation, interaction and direct contact, Fig. 3C, see methods). The duration of the event start epoch varied based on the behavior of the implanted rat and the stimulus (Fig. 3D). Using this intrinsic variability, we observed that panresponsive cells showed a brief transient period of activity at both the presentation and removal of the stimulus regardless of the stimulus identity (Fig. 3E-F, S6, S7), while the largest increase in firing in both event-specific and panresponsive neurons occurs seconds prior to direct contact with the stimulus. This increased activity likely reflects awareness of stimulus identity or the decision to interact (Fig. 3G-I). Unit activity profiles showed similar temporal patterns across the 4 classes of stimuli despite their very different sensory natures and across two days of recording (Fig. S6) strongly suggesting identification of the event and not its specific sensory characteristics.

**Fig. 3.**
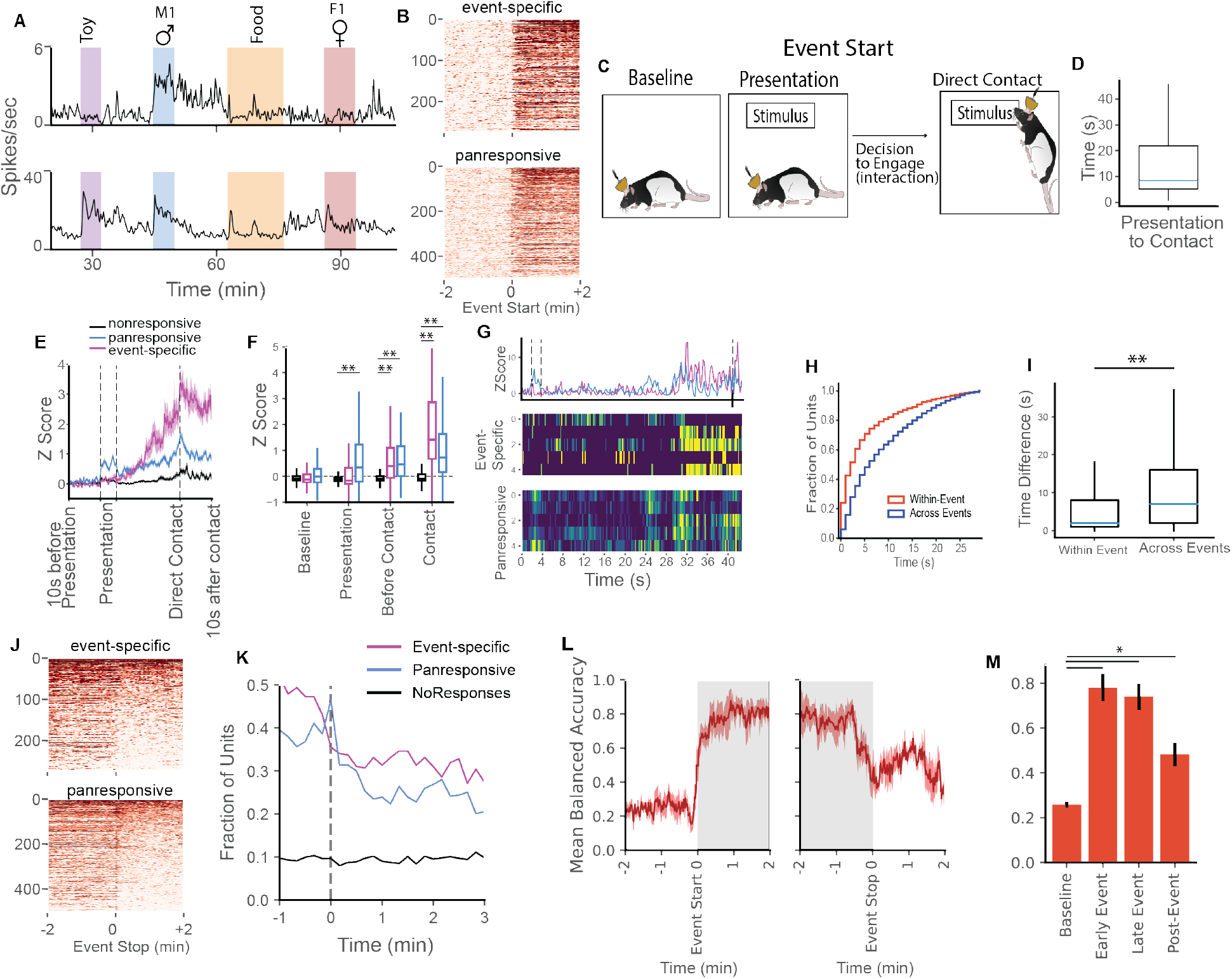
BLA neurons fire at the beginning of, during, and after an ethological event. (**A**) Abrupt increase in activity at event onset and its long slow decay afterwards in a typical (in this case male) event-specific (top) and a panresponsive (bottom) neuron. (B) Across different events, both event-specific and panresponsive neurons show sharp increases in activity with sustained activity for several minutes following event start. (C) Event start can be further divided into distinct periods based on behavior. Following baseline recording, the stimulus is placed at the opposite end of the chamber (presentation). A decision to engage is made by the implanted rat (80% of cases, i.e. active approach) or the stimulus (20% of cases, approach by conspecific) following by direct physical contact between the implanted rat and the stimulus. There is a large variation in time to contact after presentation with an average of about 10s. (D). E) Warping each event start into a common timeframe reveals differences between panresponsive and event-specific neurons, with increased activity occurring during the presentation period exclusively in panresponsive neurons and event-related firing starting in both event-specific and panresponsive before contact. This effect is quantified in (F), (**p<0.001 2-way ANOVA with multiple comparisons, event-specific and panresponsive neurons compared to nonresponsive). The step-like increases in population activity prior to direct contact in E are caused by clusters of neurons within a single event rapidly switching between an off and on state as seen in the representative example (G). Within a single event trial, individual neurons initiate their firing within seconds of each other indicating a uniform population response to the eliciting event. This is quantified (H and I) by comparing the relative timing of firing activity of responsive neurons within individual events to the relative timing of firing activity of the same neuron across different events (see methods, **p<0.01 Kolmogorov-Smirnov test, N=3231, 2026 pairs of neurons within the same event, the same neurons across different events, respectively). (J) In contrast to the rapid onset of activity at event start, after event offset many neurons continue to show elevated firing activity for several minutes (aftereffects). (K) This elevated activity occurs in approximately 30% of event-specific and panresponsive neurons and stably persists long after event stop. (L) Stimulus identity can be reliably decoded from neuronal activity both during the event (area shaded grey) and after the stimulus is removed. Decoding accuracy is significantly higher than baseline (M), (Repeated-Measures one-way ANOVA *p<0.05, N = average accuracy for each time period per 4 rats)

### Firing persists in a subset of BLA neurons after stimulus removal

In contrast to the sharp increases in firing activity seen at event onset, elevated firing continued in approximately 30% of responsive neurons after the removal of the stimulus (33 % of event-specific neurons and 27 % panresponsive compared to 9% nonresponsive, Fig. 3J-K). Fig 3A shows examples of this after-response in a male-responsive cell (top) and a panresponsive cell (bottom, see also Figure1, top 2 panels). The period of persistence differed between cells and even within cells during different presentations of the same event with some elevated firings lasting the entire post-event and others decaying at a faster rate. The ceiling effect in firing persistence in many cells precluded accurate estimates of population time constants although we have observed persistence for as long as 22 minutes in unpublished studies. All stimuli were capable of producing aftereffects and aftereffects were present on the second day of recording in slightly fewer neurons (S7). Compared to baseline, the population firing activity in the post-event period was capable of decoding stimulus identity suggesting that this activity may function as a memory trace (Fig. 3L-M).

### Firing rates of activated neurons are modulated by specific social interactions

Once firing activity was triggered by an event, specific behavioral interactions caused short-term fluctuations in firing rates. To study these micro-behaviors, we classified key social behaviors using an SVC classifier trained on manually annotated video data (Fig. S8 and Movie S5). Then, using the automatically classified behavior, we calculated the interindividual distance that best discriminated behavioral interactions from non-interactions and applied this to non-social interactions as well (Fig. S9 and see methods). The firing rates of many responsive units were modulated by active interactions with the target stimulus (Fig. 4A-C, S9). Event-specific neurons only showed micro-behavioral modulation in response to their own tuned stimulus (Fig. S9). The micro-behavioral correlation was higher at the population level than at the single unit level and there was no difference in anatomical location between units that showed micro-behavioral modulation and those that did not. (Fig. S9). 5-fold cross-validation and LDA decoding allowed reliable decoding of micro-behavioral modulation in all 4 event types (Fig. 4D) and decoding performance was strongly dependent on the number of neurons used (Fig. S10). We further examined social events to see whether single units fired in response to selected micro-behaviors including sensory behaviors (head-to-head contact, head-to-tail contact), movement related behaviors (approach and following) and passive contact initiated by the conspecific. We found units showing increases or decreases in response to specific behaviors, with head-to-head contact particularly well represented (Fig. 4E–4I, S11). Though these neurons reliably responded to specific behaviors, their responses were not unique to these behaviors (Fig. S11).

**Fig. 4.**
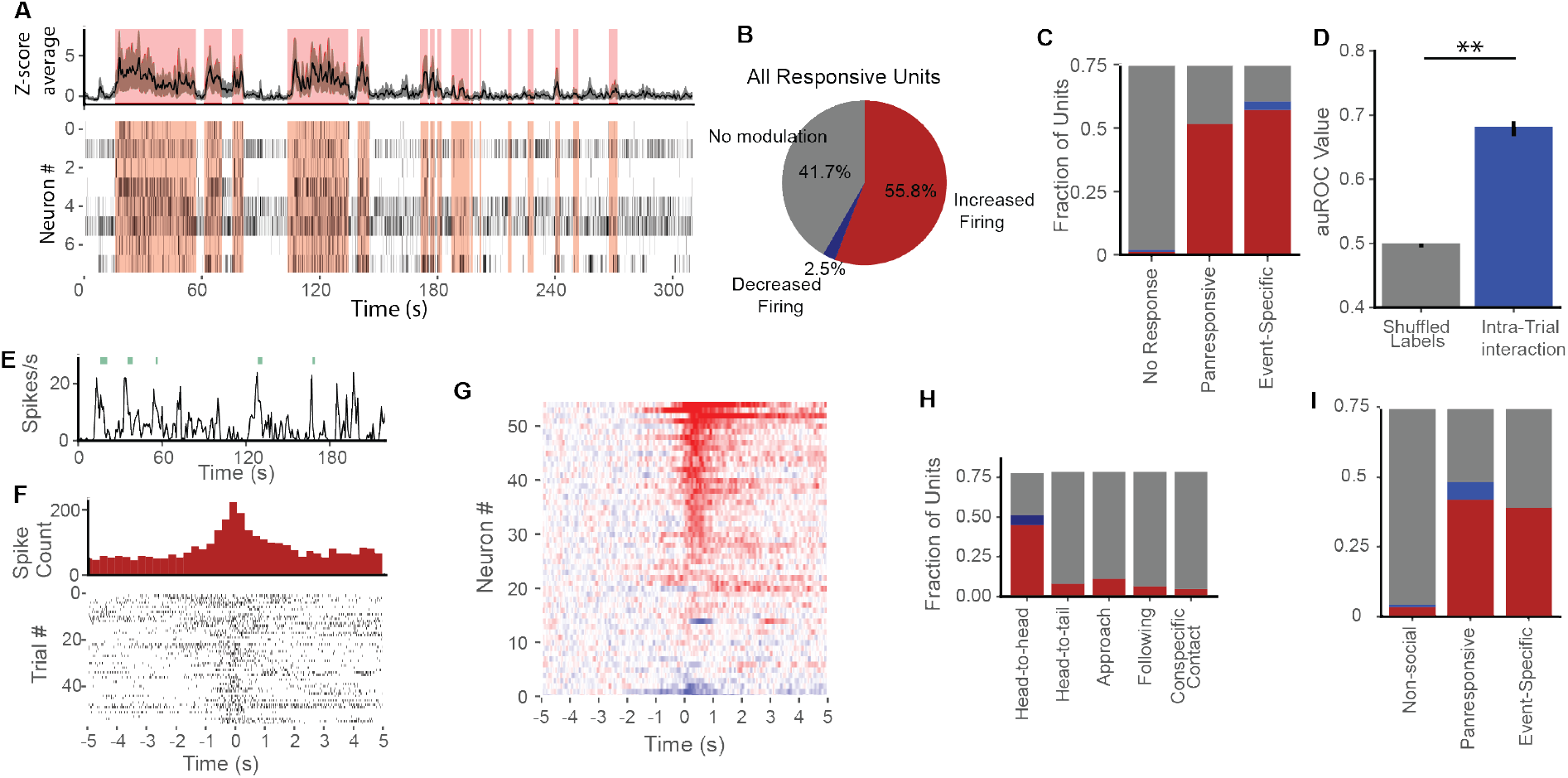
BLA neuronal activity is modulated by interaction with the stimulus with a subset of neurons reacting to discrete social behaviors. (**A**) Representative trial showing single units (raster plots on bottom) modulated by interaction with a female conspecific (social behavior shaded in pink). (B) The majority of BLA units are modulated by interaction with no differences in the proportion of panresponsive or event-specific units modulated (C)**. (**D) Behavioral interaction with a stimulus can be decoded from neuronal activity (**p<0.01 paired Student’s t-test, n=60 events). (E) Representative example of a single unit showing time-locked firing increases during head-to-head contact (green ticks). Note that the unit also bursts outside these contacts. (F) Peri-event histogram of the unit in E showing all instances of head-to-head contact (at time 0). (G) Z-scored PSTH traces for all social units sorted from greatest to least response to head-to-head contact. (H) The proportion of units that increase (red) and decrease(blue) firing to specific social behaviors by behavior type and (I) neuronal class.

### Event-specific neurons drive firing in panresponsive neurons

We identified the directionality of firing and connection strength from significantly cross-correlated BLA neurons throughout the entire recording session (see methods, Fig. 5A, 5H-J). This revealed a clear circuit structure where, typically, several event-specific neurons conveyed information to single panresponsive neurons through putative monosynaptic inputs (Fig. 5B-F, S12). Connected neurons tended to lie within 200 μm of each other (Fig. 5G). To test how these connections changed in response to presentation of the ethologically relevant stimulus, we calculated the connection strength from correlated neurons before, during and after the first presentation of each stimulus type (Fig. 5K-M). Connection strength significantly increased during the post-event period both at the population level and within continuously active pairs of correlated neurons (Fig. 5N, 5O). These results support a model where firing from event-specific neurons drives activity in panresponsive neurons, likely driving increased plasticity within the BLA.

**Fig. 5.**
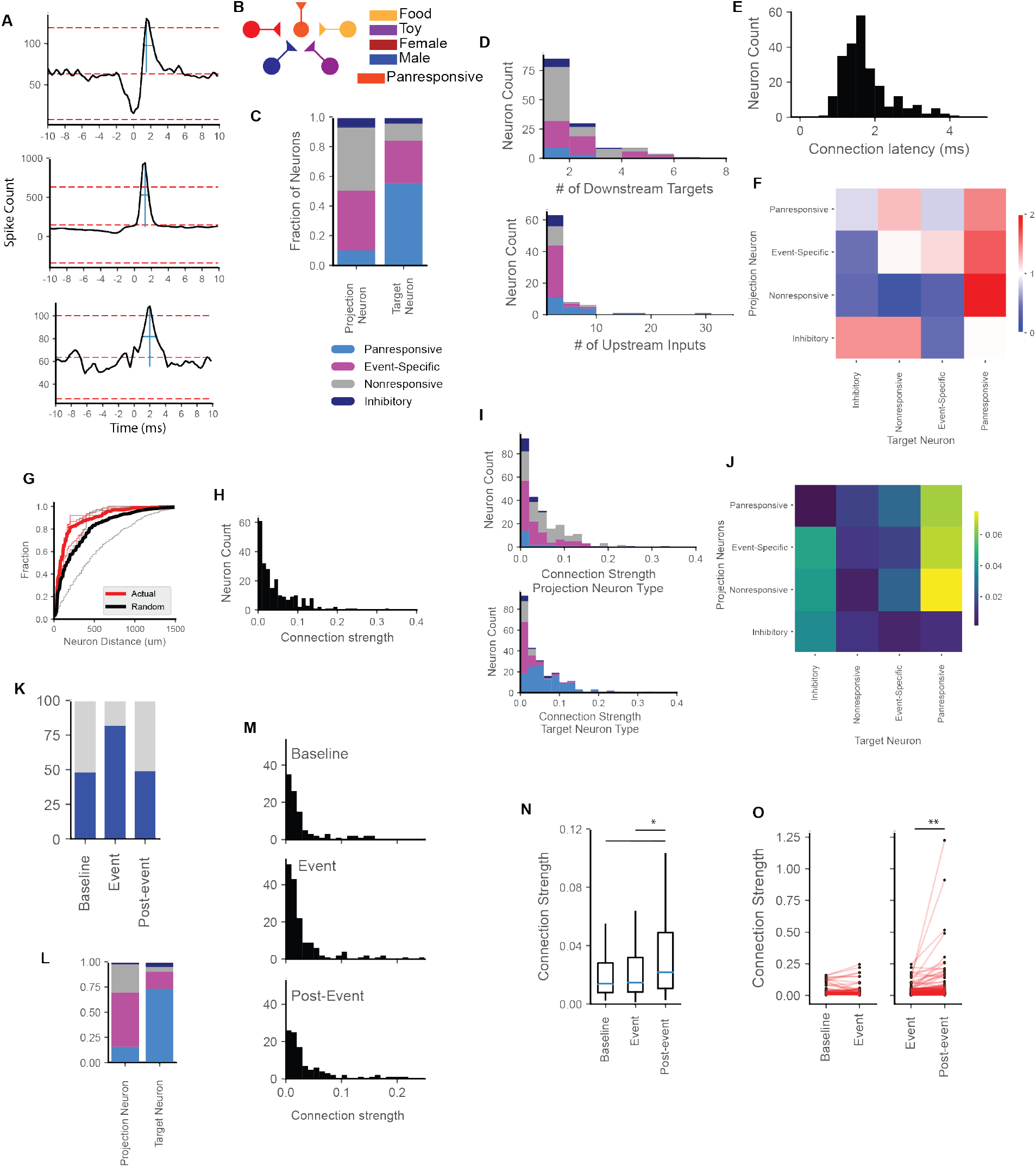
Event-specific neurons drive firing activity in panresponsive neurons. (**A**) The BLA is highly interconnected. Representative cross-correlograms show putative monosynaptic connectivity between BLA neurons. Connected target neurons exhibited a sharp increase in firing within milliseconds of the projection neuron firing (see methods). (B) BLA circuit model suggested by the cross-correlation results where different event-specific neurons drive firing in panresponsive neurons. (C) Throughout the entire recording session, we identified 234 cross-correlated pairs of BLA neurons. Within these pairings, the majority of projection neurons were event-specific or nonresponsive, while the majority of target neurons were panresponsive. (D) Individual projection neurons had few downstream targets (top), while target neurons often received input from many projection neurons. (E) The majority of correlated neuron pairs fired within 1-2ms of each other suggesting putative monosynaptic connections. (F) Connectivity between specific types of neurons was seen more often in the BLA than expected by chance (>1 = more likely than chance, <1 = less likely than chance). (G) The majority of connected neuron pairs were located within 200um of each other, closer than expected by chance. (H) We calculated connection strength between pairs of neurons. Panresponsive neurons were more likely to receive stronger input from event-specific, nonresponsive and other panresponsive neurons (I-J). (K) We identified the number of neuron pairs that showed significant cross-correlations before, during and after the first presentation of each stimulus. L) Similar to the crosscorrelations seen throughout the entire recording, projection neurons were more likely to be event-specific or nonresponsive neurons whereas target neurons were predominantly panresponsive. M) Connection strength increases in the post-event period. This is quantified in (N) (*p<0.05, Kruskal-Wallis test with Dunn’s correction, N = correlated pairs N=100, 157, 107 for baseline, event and post-event period. O) Individual neuron pairs increased in connection strength between the event and post-event period (p = 0.54, baseline-event connectivity change, N= 65 neuron pairs; **p<0.01, Student’s paired t-test, event-post-event connectivity change, N=157 neuron pairs).

## Discussion

The BLA contains information about many important aspects of the ethological events studied in this experiment. Unlike other areas often described as part of the social circuitry (e.g. mPFC, MEA) (*14*, *15*), the BLA as a whole did not show a strong preference for social or non-social events. Instead, such a preference was manifested at the level of BLA subdivisions with nonsocial (food and toy) events represented in the LA and BA and social conspecific events in the ventral BA and BMA. BMA, and to a slightly lesser extent, BA further distinguished strongly between males and females. Subnuclear preferences for different events are consistent with reported anatomical differences in BLA connectivity(*16*). The similarity in temporal response profiles across events and anatomical subdivisions suggests that despite the recruitment of different neuronal clusters, the coding principles used by BLA neurons (Fig. S13) during different ethological events are likely similar at the cellular and circuit level.

At the single-cell level, we identified two main classes of BLA neurons. Approximately 30% of BLA neurons (event-specific neurons) exhibited highly-tuned macro-scale increases in firing to one of the four ethological events in our study. These neurons were spatially localized, encoded the identity of the ethological stimuli and continued to signal the most recent event for minutes after its termination. These neurons drove firing in a second class of putative BLA interneurons (panresponsive neurons), which fired in response to multiple ethological events, signaled the beginning and the end of each event by brief bursts of action potentials, and received input from multiple neurons including event-specific neurons and other nonresponsive neurons. Given the stability of individual event-specific neuron’s responses to a single ethological stimulus in our study, we suggest that these BLA neurons may be hard-wired to respond to innate naturalistic stimuli (for example male or female conspecifics or food), and hypothesize that the detection of these stimuli is one of their primary functions, reminiscent of neuronal responses to salient stimuli like faces previously reported in other brain areas(*17*). On this view we would predict that many of the neurons in the BLA classified as nonresponsive would respond selectively to other ethologically significant stimuli such as rat pups, nests, other salient locations, prey or predators. In our study, nonresponsive neurons resembled event-specific neurons with low baseline firing rates, broad waveforms and similar connectivity to panresponsive neurons. We further speculate that continued experience with different males, females or types of food might lead these cells to learn to distinguish individual animals or foods, respectively.

Once activated by a given event, the firing patterns of many BLA cells are modulated by the microstructure of the event, most notably by active interaction with the stimulus. This suggests that activated clusters of BLA neurons represent finer grained details of the event similar to the state-changes described in BLA calcium imaging studies (*4*, *5*, *18*),including specific sensory social behaviors, most predominantly head-to-head contact. Our study looked only at a limited set of specific behaviors associated with sensory inputs, and it is likely that BLA neurons are modulated by a host of sensory and behavioral factors (e.g. smells, taste, auditory vocalizations, etc.). For example, it is known that social transmission of olfactory information about edible foods is transmitted by the breath of the informing animal and is amygdala-dependent(*19*). However, unlike the large-scale gating of activity that occurs in single-units during events, these responses were non-specific and less robust at the single-cell than at the population level.

In addition to providing evidence that the BLA is involved in identification of ethologically salient stimuli, the results show that it continues to provide information about the specific event for minutes after its cessation. We suggest that this could form the basis for an active memory trace of the event. The BLA has a well-established role in memory consolidation and recall(*20*), and inhibition of the BLA or activation of specific ‘engram’ cells can affect this process(*21*, *22*). The aftereffects seen in our data might support short-term active memories allowing subsequent recognition and recall of the event or might also be required for consolidation in other brain areas or the elicitation of long-term hormonal responses by these events. These effects have been observed at smaller scales in other brain regions (e.g. cortex, hypothalamus) (*23*)(*24*), but the BLA responses are unique in their long time scales often exceeding 5min in duration. One argument against a nonspecific consolidation theory is the specificity of the trace for the actual event experienced. We hypothesize that this long-term persistent firing seen in the BLA likely originates within the BLA circuit, with event-specific neurons triggering firing in BLA interneurons, which may release neuropeptides or other signaling molecules that maintain high activity in the BLA for long periods of time. Additionally, BLA projection neurons likely drive activity in higher-cognitive areas (e.g. mPFC, hippocampus) that are associated with memory formation. It remains to be seen how BLA projection neurons relate to the functional classes described in this study (e.g. do the same event-specific neurons that send local BLA connections to panresponsive neurons also project to other brain structures?).

Overall, the results suggest that the basolateral amygdala is involved in the detection of ethologically significant stimuli, the signaling and perhaps generation of appropriate behavior towards those stimuli, and the continued neural representation of the event in the form of an active memory for considerable periods of time after its occurrence.

## Acknowledgments

We thank Yoh Isogai and Daniel Regester for design of the retrievable Neuropixels holder and members of the O’Keefe lab for helpful discussions regarding the results and manuscript.

## Funding

This project has received funding from the European Union’s Horizon 2020 research and innovation programme under the Marie Sklodowska-Curie grant agreement No. 840562 (CM).

Sainsbury Wellcome Centre Core Grant from the Gatsby Charitable Foundation and Wellcome Trust (090843/F/09/Z, JOK)

Wellcome Trust Principal Research Fellowship (Wt203020/z/16/z, JOK)

## Author contributions

Conceptualization: CM, JOK

Analysis/Investigation: CM

Funding: CM, JOK

Writing: CM, JOK

## Competing interests

Authors declare they have no competing interests.

## Data and materials availability

All data and code will be made available upon reasonable request to CM or JOK

## Notes

### Competing Interest Statement

The authors have declared no competing interest.

