## supplemental information for "Basolateral Amygdala Represents and Remembers Ethological Events"

**Materials and Methods**
Animals
All animal procedures were conducted in accordance with the UK Animals (Scientific Procedures) Act (1986). All rats were housed in large clear plastic cages and maintained on a 12h reverse light/dark cycle in a temperature and humidity-controlled room. All rats used in these experiments were adult male or female Lister Hooded rats.
The 5 adult male rats which received Neuropixels implantsranged between 450-600g, were 3-6 months of age, and had never been housed or exposed to female conspecifics prior to the experiment. Prior to surgery, they were housed 2-3 per cage and single-housed after Neuropixels implantation. Implanted rats were fed once per day and maintained at least 90% of their pre-restriction body weight. Both female and male rats were used as social conspecifics. Females were between 4-12 months of age to ensure possible sexual receptivity and male conspecifics were between 3-6 months. Conspecific rats were housed 2-3 per cage and received ad libitum access to food. All rats (conspecific and implanted) received ad libitum access to water.

Neuropixels Implantation
To record from BLA neurons, rats underwent stereotactic surgery to implant a Phase 3A Neuropixels probe (*25*) targeting the left BLA. Rats were anaesthetized with isoflurane in $O_2$ (2-3%) and given pre-op analgesia subcutaneously (Carprieve 5mg/kg) before being placed in a stereotaxic device. All 5 rats were implanted in the left hemisphere (BLA coordinates relative to bregma, -2.8mm anterior, 5mm lateral and -9mm ventral). A stainless steel reference screw located in the right frontal cortex was connected to the ground/reference cable of the neuropixel probe during surgery. 2-4 additional screws were threaded into the skull to support the anchoring of the implant with superbond and light curable epoxy. The implant was protected with a copper mesh cage that provided physical protection to the probe, being grounded to the probe/reference screw, shielded electrical noise . The opening of the copper mesh cage was wrapped with 3M Coban wrap when the probe was not in use.
4 rats received retrievable neuropixel implants where the neuropixel probe was dipped into concentrated Di-O (Neuro-DiO, Biotium, 50mM in isopropanol) 1-2 h prior to implantation. Briefly, the Neuropixels probe was epoxied onto a 3-part CNC machined holder that permitted retrieval of the intact probe post-experiment for later re-use. 1 rat received a neuropixel probe permanently implanted in a custom-designed 3D printed holder. Post-surgery rats received analgesic and antibiotic mixed in strawberry jelly for 3 and 5 days respectively (Metacam 1mg/kg, Baytril 1%). Animals were allowed to recover for 48 h before a test baseline recording and 72 h before full experiments commenced.
Neuropixel probes were retrieved for reuse after the experiment was completed in 4 rats (average duration of recordings was 7-10 days). Briefly, the rat was reanesthetized (isofluramce in O2, 2-3%) and the copper mesh cage surrounding the implant was removed. Since the neuropixel probe was solely attached to 1 part of the CNC machined holder, we carefully unscrewed this part from the rest of the holder that was affixed to the skull with superbond. The probes were carefully removed, tested, cleaned (1h in 1% Terg-a-zyme in MilliQ, Sigma-Aldrich, followed by 1h in milliQ) and reused (3 probes used in 4 rats). Anaesthetized rats immediately received an overdose of sodium pentobarbital prior to perfusion.

Histology
After completion of the recording session and/or probe retrieval, rats under isoflurane anaesthesia received an overdose of sodium pentobarbital and were transcardially perfused with phosphate-buffered saline (PBS) and 4% paraformaldehyde (PFA). In the 4 rats that received a Di-O dipped Neuropixels probe , brains were transferred to phosphate buffer (50mM PB) and then coronally sectioned in 50um steps while simultaneously imaged using a custom serial two-photon tomography microscope. For the other rat, the brain was transferred to 30% sucrose in PBS for at least 48 h and frozen on dry ice prior to sectioning on a cryostat. Frozen coronal sections cut at 40 um were collected and stained for GFAP expression (primary antibody: DAKO anti-rabbit GFAP 1:500 in PBSGT - 1% normal goat serum and 0.3% Triton-X in PBS; second antibody: goat anti-rabbit Alexa 568 1:500 in PBSGT). Mounted sections were imaged using a Zeiss Axioscan. Regions of the neuropixel probe within the BLA were determined by manually registering the Di-O or GFAP track onto the Paxinos brain atlas and mapping the coordinate channel locations of the neuropixel probe onto the track.

Data Collection/Experimental Setup
Neuropixel recordings took place in a custom-made rectangular acrylic box measuring 100cm x 70cm with 45cm high walls. The chamber was illuminated by 6 LED photo lights (F+V K320 Lumic Daylight LED Video Light,

Wex lighting) for consistent illumination of the space. Continuous video recordings (minimum 30Hz frame rate) were taken with a Basler color video camera (acA2500-60uc Basler Ace USB Camera with Kowa LM8HC camera lens) focused to the floor of the chamber. Neuropixels recordings collected at 30KHz were multiplexed and digitized on the probe and sent via a connected cable to an FPGA board. Extracellular recordings were collected and saved in binary format using SpikeGLX software (https://github.com/billkarsh/SpikeGLX). The video camera was controlled through custom-written software on labview which saved compressed avi video files of the experimental events and sent a 5V synchronizing signal to the Neuropixels board (National Instruments 2018 with NI USB-6000 for 5V pulse generation). To stabilize the Neuropixels cable and headstage and prevent pulling of the delicate neuropixel flex cable, we 3D printed a custom-designed holder that held and protected the headstage and cable connection. This was secured to the copper mesh cage on the rat's head using alligator clips and counterbalanced with a small weight. After a baseline recording and habituation to the recording chamber, we started experiments. Rats implanted with Neuropixels probes were connected to the recording apparatus and allowed to freely explore the chamber. The counterbalance was adjusted as needed for the individual rat. During a single experimental day, rats were presented with a battery of different presentations including female conspecifics, male conspecifics, remote-controlled toy mice (HEXBUG 480-4466-00TG12 Remote Control Mouse Cat Toy) and sweetened cooked rice. Following 5 min of baseline recording, each stimulus was placed into the box on the opposite side of the implanted rat and the rat and stimulus were allowed to freely interact. Social interaction was only discouraged when there was danger to the Neuropixels probe (i.e. conspecific rat climbing upon or attempting to chew on the exposed bit of the probe). After 5 min of free interaction with the stimulus, the stimulus was removed from the box during a suitable window (i.e. when there was a break in the social interaction or in the case of food when the rat had finished eating). Recordings were continued for an additional 5 min prior to the chamber being wiped clean of any feces or urine and the start of the next event sequence. In a randomized order, social conspecifics were presented 2-4 times each while non-social toy and food were presented once each. The entire battery of presentations was then repeated once more in the same order. Rarely, the implanted rat had to be carefully untwisted due to excessive turning in one direction. To do so, between different stimulus presentations, rats were carefully lifted and untwisted to preserve recording integrity. Experiments were repeated for a minimum of 2 days. 1 of the 5 rats did not receive repeat presentations of stimuli and the ability to track multiple conspecifics in an environment was lacking, therefore the data from that rat was only included in Figures 1A-G and 2A-C.

Single Cell Isolation from Multi-unit recordings

Neuropixels and video data were saved in a single file each for individual events. Neuropixels data was concatenated across an entire recording day, noisy channels were removed from analysis and the rest of the data was median subtracted to remove artifacts common across the probe. Single units were isolated through Kilosort 1.0 (https://github.com/cortex-lab/KiloSort) and manually curated using Phy or Phy2.0 (https://github.com/cortex-lab/phy). Single units were manually curated according to the following criteria: less than 0.1% of spikes violated the cell refractory period of 2ms, spike waveform was consistent with a single unit, amplitude of at least 40mV and absence of any 50Hz noise in autocorrelogram. Waveform and crosscorrelograms of all nearby units were compared to verify that there were no two clusters corresponding to the same neuron.

Synchronization of Video and Electrophysiology

During each frame acquisition of video data, the labview program gave a random 80% chance that a 5V pulse would be sent out to the Neuropixels board. The resulting pulse or no pulse was recording by the program along with the relative timing, preserving a copy of the exact synchronizing spike train that represents the camera frames. We extracted the received spike train from the synchronizing channels of the Neuropixels board through a custom-written python program and verified that the pulses received by the Neuropixels board were of the exact number and sequence as the pulses recorded sent by Labview. We extracted the timing of the received pulses by the Neuropixels board, interpolated based on framerate for the 20% of not-sent pulses and used this as the synchronizing index for synchronizing video to electrophysiology.

Tracking from Video Data

Prior to experiment, the white fur on the tailbase of 4 implanted rats was marked with a purple marker. Conspecific rats also received fur colorings on their right ear, left ear and tailbase with yellow, orange and blue colors, respectively. All conspecific rats received the same color patterns. This color pattern was replicated on the moving toy mouse to approximate the 'ears' and tailbase.

All video data was post-processed offline to identify the coordinate positions of all animals and objects in the recording chamber. We trained Deeplabcut 2.0.1 (*26*) on videos from 3 different recording sessions which contained different implanted rats and different conspecific rats. Using this trained network, we were able to reliably identify 10 different body parts per rat across all videos. On the implanted rat we identified left and right headstage, head location, left nape, right nape, center nape, left mid-back, right mid-back, center mid-back and tailbase. On conspecific rats we identified the left ear, right ear, nose, left nape, right nape, center nape, left mid-back, right mid-back, center mid-back and tailbase. Deeplabcut tracking of the experimenter-colored body parts had better accuracy, so we thresholded the other body parts per rat to exclude data points where there was an unnatural distances between body parts. We also excluded all points that were under 0.999 probability for the experimenter-colored body parts and 0.9 for the other body parts. Data was then interpolated linearly to fill in gaps.

From the xy coordinates of the 20 tracked body parts, we calculated features relevant to social behavior. The calculated features are as follows: calculated independently for the implanted rat and conspecific; head velocity, tailbase velocity, spine length, head angular velocity, and tailbase velocity offset by 1s; from both animals: interindividual distance between different body parts (head, tailbase, nape, and back), headdirection from one rat to the other rats different body parts, the difference in head angle between the implanted rat and conspecific, the difference in interindividual distance between the rats' heads and the implanted rat's head and conspecific's tail, the difference in the headdirection between the rat's heads and the implanted rat's head and conspecific's tail, and the correlation between head and tail positions over 1s. We removed outliers from these features and interpolated any missing data.

Classifying responsivity of individual neurons

We evaluated the response of each individual neuron to each event or presentation of a specific stimulus. Briefly, we calculated the firing rate of each individual neuron with 1s bins and compared the firing rate from the 5min event period to the firing rate of the immediately preceding 5min baseline period. We computed the receiver operating curve (ROC) and from that calculated the area under the receiver operating curve (auROC) and converted this into a response score (auROC – 0.5)*2. We assigned a response score of 0 (auROC of 0.5) to neurons that had low firing rates during the baseline and event periods (less than 50 spikes).

To calculate the average response score, we averaged the response score for each of the 4 stimulus types (male, female, food, and object) and set the threshold for responsiveness at +0.2. Neurons that had a response score above 0.2 for more than one modality were classified as multimodal and split into 2 categories – panresponsive (significant scores on both social and non-social stimuli) or non-social/social. A large proportion of units exceeded the threshold on only one specific stimulus type (event-specific) and these were classified based on their stimulus preference. Units that did not increase firing in response to any stimulus presentation were further split into 2 groups (non-responsive and firing decreases). Neurons that decreased firing had average response scores less than -0.2 for at least one modality.

Classification of Putative Pyramidal Neurons and Putative Interneurons

We calculated the waveform for each recorded neuron by extracting the raw voltage trace from 500 randomly selected spiketimes per neuron. We averaged these voltages together and calculated the time delay between the trough and peak. A small fraction of waveforms had inverted waveforms (where the peak voltage precedes the trough) and these were excluded from the analysis and classified as irregular waveforms. To divide the rest of the neurons into putative pyramidal or interneurons we plotted the trough to peak distance against the baseline firing frequency from the first 5 min of each recording day. We divided the results into 2 groups using K-Means clustering, with putative pyramidal neurons having longer trough to peak distances and lower baseline firing rates and putative interneurons having shorter trough to peak distances and higher firing rates.

Population Correlation of Response Scores

To calculate the correlation between population response scores across different stimulus presentation, we calculated the population vector for all responsive neurons (multimodal and event-specific) for each stimulus presentation. To combine population vectors across different rats, we defined social presentations (e.g. female1 and female2) as being the first female conspecific presented and the second. If there was a third female or male conspecific presented to that rat, they were not shown in Figure 1H, but included in the quantification in Figure 1I. We calculated the Pearson's correlation between the calculated population vectors.

Decoding Stimulus Identity from Neural Activity

To test whether the population response score accurately represents the stimulus identity, we used Linear Discriminant Analysis (LDA, python package Sci-kit learn) (Figure1). For each rat, we trained the LDA decoder on all the population response scores of all the stimulus presentations except one and then tested on the omitted dataset

(leave-one-out method). We repeated this until we had a predicted stimulus type of each stimulus presentation and calculated the accuracy of these labels against the correct labels, giving one score per rat. We then repeated 500x with shuffled labels and calculated the control score per rat.
To calculate how the number of neurons used in the decoder affected performance (S4), we trained the LDA decoder on a random selection of neurons at increasing numbers (from 2 neurons to the maximum number of neurons per rat, 500x per neuron number selection). We then repeated this analysis, but instead of using all neurons in the dataset we omitted specific classes of neurons (panresponsive, event-specific and nonresponsive). We calculated the change in performance of the decoder for each rat by subtracting the performance using all neurons from the performance with the omitted neurons controlling for neuron number.
To calculate how the location of neurons affect the accuracy of the LDA decoder for social events (Figure2), we recalculated the accuracy using the leave-one-out method for all social events but only included neurons from the same rat located within 500um of each other, with a minimum of at least 10 neurons included in the dataset.
To calculate how firing around the time of stimulus contact affected the performance of the LDA decoder, we used the leave-one-out method described above to calculate the accuracy of an LDA decoder trained on a shifting window of 10s of neuronal firing data from 30s before stimulus contact to 30s after contact. We compared the accuracy per rat in 5s epochs from 10s before stimulus contact to 5s after.

Quantification of differences in firing onset and firing offset
We calculated firing rate data at event start with bins of 100ms per neuron. We z-scored the firing rate to the baseline firing rate per event (firing rate – mean baseline firing rate / standard deviation baseline firing rate). For visualization across different events (Figure 3E, S6), we time-warped individual z-scored traces using Spline Interpolation into a common timeframe.  All quantification was performed on the non-interpolated raw z-scored data (Figure 3F).
To calculate whether individual units within a specific trial started firing at similar times (Figure 3G-3I), we calculated the time delay to peak firing per unit after stimulus presentation but before direct contact. We calculated the Euclidian distance between these values for all units within a single trial. We compared this to the Euclidian distance of the time delays between the firing onsets of the same unit within different trials.
To calculate aftereffects (Figure 3J-K), we once again took z-scored firing rate data from individual stimulus presentations and sorted based on largest average z-score difference in the post-stimulus presentation period relative to baseline. We calculated the proportion of neurons with an average z-score above 1 in 10s bins for 3 minutes after stimulus removal.
To test whether neural activity before and after interaction with a stimulus can decode stimulus identity, we trained an LDA classifier on response scores during the event period. We calculated the response score from 2 minutes before event start to 2 minutes after event stop in 10s bins and used these bins as the test data.

Automatic Classification of Behavior from Video Data
To facilitate the analysis of whether neurons respond to specific social behaviors, we trained a support-vector classifier on 7 videos of manually annotated social behavior. We annotated videos frame-by-frame with one of 7 different labels (non-social, head-head, head-tail, approach, following, conspecific-initiated contact and other social) as detailed.
Non-social: No social engagement from either the implanted animal or conspecific.
Head-to-head: Implanted animal and conspecific facing each other and making direct contact with snout or whiskers
Head-to-tail: Implanted rat less than one head's distance from the ano-genital region of the conspecific
Approach: Implanted rat moving towards a stationary conspecific
Following: Implanted rat greater than one head's distance from the conspecific with both animals facing and moving in the same direction
Conspecific-initiated contact: Implanted rat is turned away from the conspecific, while the conspecific is engaged with and touching the implanted rat (usually sniffing the implanted rat's backside or flank).
Other Social: All other social engagement, which includes but is not limited to social grooming, boxing, flank nipping, mounting.
Using the trained SVC classifier, we were able to automatically and reliably identify social and non-social periods as well as specific social behaviors including head-to-head contact, head-to-tail contact, approach, following and conspecific-initiated contact. We used 30 features calculated from the xy coordinates for the 20 body parts on the 2 different rats and trained the support vector classifier with rbf kernel. Using 5-fold cross-validation, initial tests revealed that the SVC classifier correctly labeled data with 97% accuracy. We retested the SVC classifier by omitting one out of 7 of the manually annotated datasets in each iteration and determined that the SVC classifier had

an accuracy of approximately 90% for frame-by-frame annotation prediction. We then trained the rest of the collected videos on the trained SVC classifier of the manually annotated data and post-processed the bouts by removing behavior gaps shorter than 0.25ms and behaviors that lasted for less than 5 frames.

We performed a more basic automatic classification of non-social videos. For interaction with a moving toy object, we calculated the optimal pixel threshold for differentiating social contact vs no-contact from the automatically identified social video data and applied this threshold to the videos depicting toy interaction. For the food interactions, given that the rat was stationary while eating the sweet rice, we used the video and tracking data to draw an ROI at the position where the rat was eating the sweet rice and used this to calculate moments of eating vs non-engagement with the sweet rice.

Calculating unit response to behavior

To calculate whether individual units increased or decreased their firing in response to specific behaviors, we compared firing rates of each neuron for periods of interaction to periods of non-interaction during the event that occurred after the point of first contact with the stimulus. We calculated the auROC and response score for the comparison of these two distributions for each individual event and calculated the average response by averaging the response score for all the events that the unit responded to (e.g. interaction score for event-specific units was calculated only from those events, etc.). Units that had an interaction score greater than +0.2 or less than -0.2 were classified as increasing or decreasing firing in response to interaction.

To calculate whether units experienced time-locked increases or decreases to specific behaviors, we extracted the timestamps of all individual behaviors (head-head, head-tail, approach, following, conspecific-contact) that lasted more than 1s. We calculated the peri-event stimulus histogram for each behavior by extracting all spike times from five seconds before to five seconds after and summed the spikes binned in 100ms. We z-scored the resulting average traces per neuron to the first 3 seconds and considering any neuron showing a z score above +5 or below -5 to be responsive to that particular behavior.

Decoding Behavior from Neural Activity

We took the z-scored population data with labels for interaction or non-interaction per individual event and calculated the ability of an LDA decoder to predict whether the animal was interacting or not interacting with the stimulus using 5-fold cross validation. Each dot represents a different event comparing the predicted score against the actual score using ROC analysis. We reran the analysis with different numbers of neurons as well as separating the events based on stimulus type.

Cross-correlation Analyses

To identify pairs of cross-correlated neurons across the entire recording session, we calculated the cross-correlogram for every simultaneously recorded neuronal pair with a resolution of 0.25 ms. To determine significance, we identified cross-correlograms with increased firing that exceeded 3 standard deviations. We used a peak detection algorithm on the cross-correlogram between -5 to +5 ms (scipy.signal.find_peaks, height = 3 standard deviations above mean, width = 0.75 ms) and filtered the dataset to only include cross-correlograms where the peak occurred after 0 ms and the peak width did not start before 0ms (representing directional putative monosynaptic connections). We excluded neuronal pairs where either the projection or target neuron fired less than 100 spikes or when the peak firing in the cross-correlogram was lower than 20 spikes. To determine neuronal connectivity during the first presentation of each stimulus type, we determined how many of the previously identified cross-correlated neurons showed significant cross-correlations during the baseline, event or post-event period for the 4 stimuli using the same criteria as above.

To determine whether connectivity between certain types of neurons was overrepresented in the dataset compared to chance, we calculated the actual incidence of different connection types in the dataset and compared that to theoretical incidence of the connection type if the neurons were randomly connected (1000x random sampling between the observed projection and target neurons).

To calculate connectivity strength, we determined the number of spikes that occurred during the detected cross-correlogram peak and divided that by the total number of spikes fired by the target neuron during the epoch. To determine whether changes in connectivity occurred within individual neuron pairs, we identified neuronal pairs that showed connectivity between two consecutive epochs (baseline and event or event and post-event) and compared their connectivity strengths.

Statistics

We used PRISM 8.0 and stats-models Python packages to calculate statistical significance. Individual statistical tests are as noted on the figures, where normality could not be assumed we corrected using the equivalent non-parametric test. P values are denoted by the number of stars, *, ** representing $p < 0.05$, $p < 0.01$, respectively.

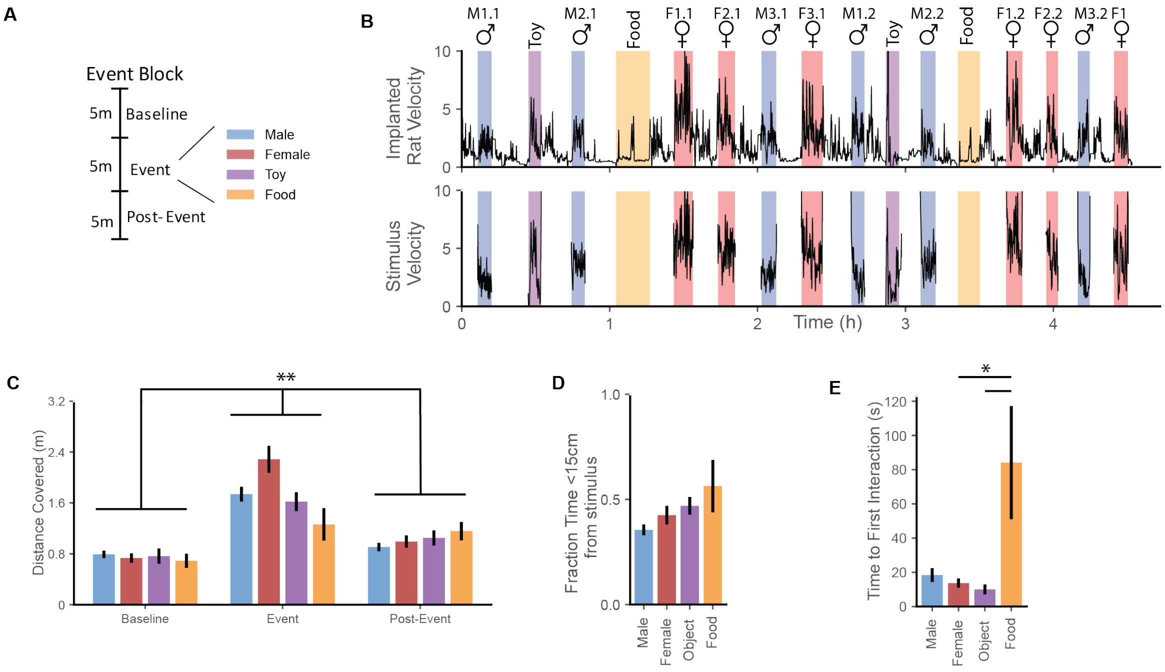


**Fig S1. Presentation of a battery of social and non-social stimuli and resulting behavior.** (**A**) Representation of a single event block and different stimuli or event-types presented. During a recording session (representative example in B), an implanted rat was presented with up to 16 different event blocks. The implanted rat was continuously tracked throughout the entire recording session and non-stationary stimuli (male conspecifics, female conspecifics, moving toy) were tracked when present. C) The total distance covered by the implanted rat increases during events with the presentation of stimuli, but there are no consistent differences between different stimuli. (**$p<0.001$ for time, $p=0.18$ for stimuli-type, repeated measures 2-way ANOVA) D) Implanted rats spend similar amounts of time within 15cm of stimuli indicating that there was not an overall preference for one type of stimuli over another. ($p = 0.17$, Brown-Forsythe and Welch ANOVA) E) Rats take slightly longer to approach and interact with food than moving stimuli (*$p<0.05$, Kruskal-Wallis test with multiple comparisons).

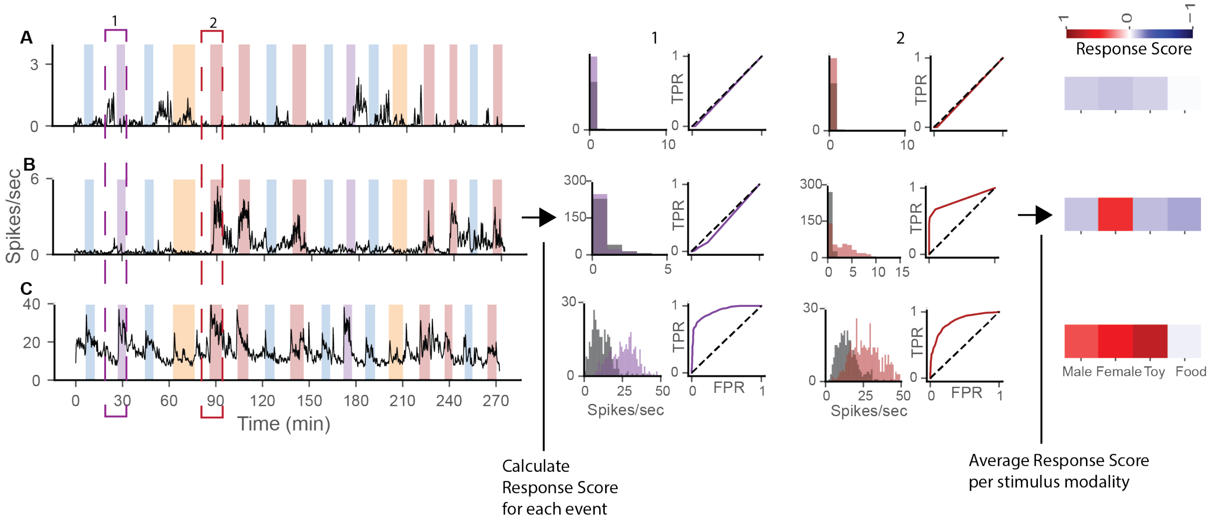


**Fig. S2. Calculation of response scores for presentations of different stimuli.**
For each neuron, we first computed the receiver operating characteristic curve (ROC curve) and calculated the area under the curve (auROC, between 0-1). We averaged together the auROCs for each event (male, female, toy, food) and converted this into a response score (between -1 and 1). Neurons were classified based on their response scores on each of the four events. A) A representative unresponsive neuron that does not have a positive response to any event. B) A representative event-specific neuron, in this case firing only to females. These neurons show a positive response score to only one event. C) A representative panresponsive neuron, which shows increases in firing to both social (male/female) and non-social (toy/food) stimuli. Dashed boxes show the periods over which the ROC was calculated for the toy and the first female events.

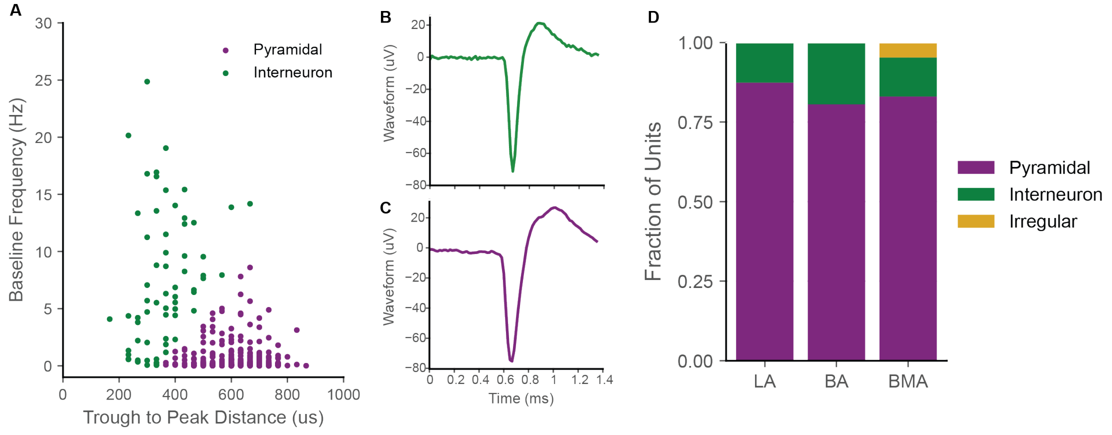


**Fig. S3. Identifying putative pyramidal and putative interneurons.**
A) Neurons were classified using K-Means clustering as putative pyramidal or putative interneurons based on their baseline firing frequency (first 5 min of the recording) and the distance between the trough and peak waveform. A representative putative interneuron is shown in (B) and a representative pyramidal neuron shown in (C). D) Small regional differences existed between different BLA nuclei with fewer interneurons being present in the LA than the BA/BMA. Irregular waveforms (sometimes called dendritic spikes, where the only waveform peak precedes the waveform trough) were only present in the BMA and were excluded from the analysis in (A).

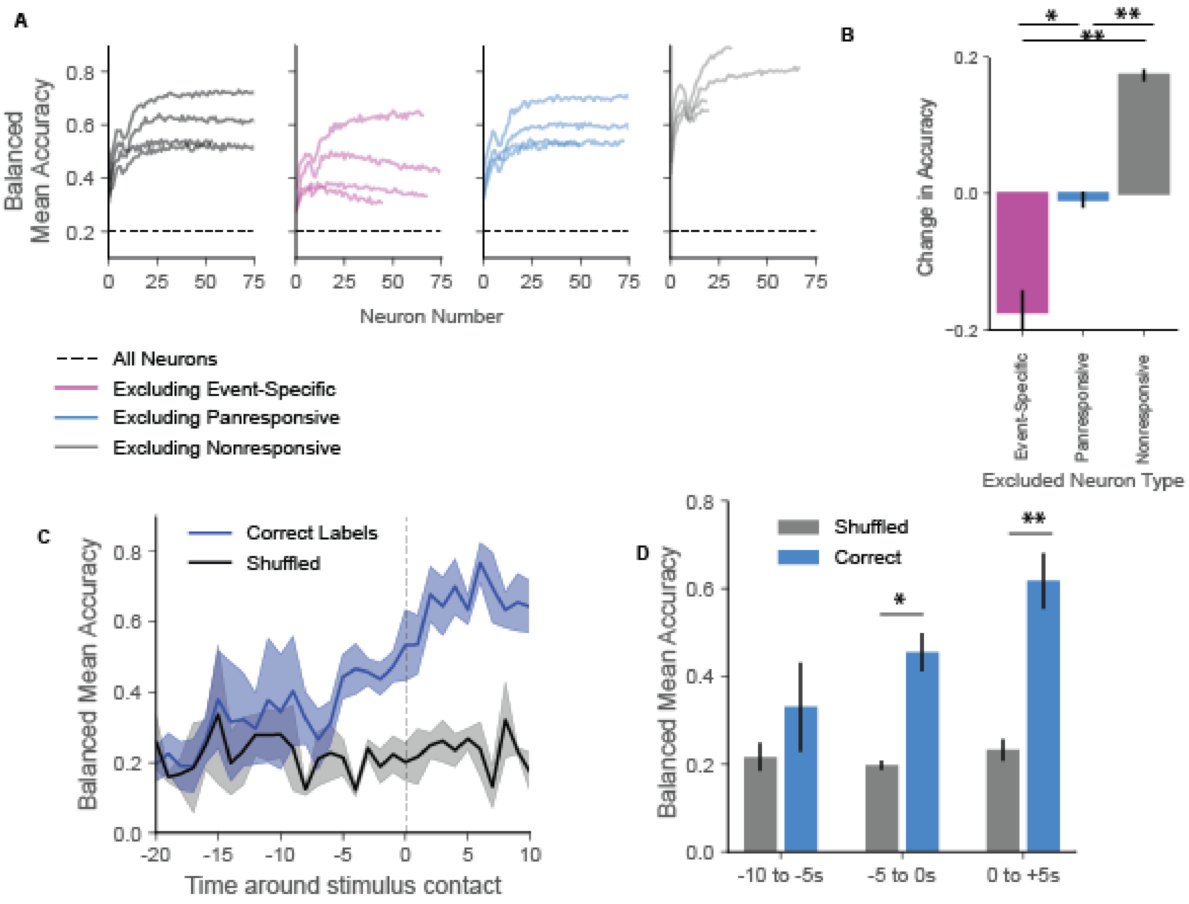


**Fig. S4. Reliable decoding of stimulus identity.**

A) Stimulus identity can be reliably decoded using response scores from relatively few neurons. By excluding specific neuron types (event-specific, panresponsive or nonresponsive) we can measure the relative contribution of each neuron class (each line represents a separate rat, see methods). B) Event-specific neurons carry more information about stimulus identity than panresponsive neurons, which in turn carry more information than nonresponsive neurons. Removing event-specific neurons impairs performance, while removing nonresponsive neurons improves performance (* $p < 0.05$, ** $p < 0.01$, N = 4 rats, repeated measures one-way ANOVA with Tukey's multiple comparisons test). C-D) To determine whether contact with a given stimuli affected decoding of stimulus identity, we reran LDA using mean firing frequency over 10s around the start of direct contact with the stimulus. Trial identify could be reliably decoded from neuronal activity starting from approximately 5s before direct contact with the stimulus. ($p=0.40$, -10 to -5s; * $p<0.05$, -5 to 0s; ** $p< 0.01$, 0 to +5s, repeated measures 2-way ANOVA with multiple comparisons, N = 4 rats, average balanced accuracy over 5s epochs)

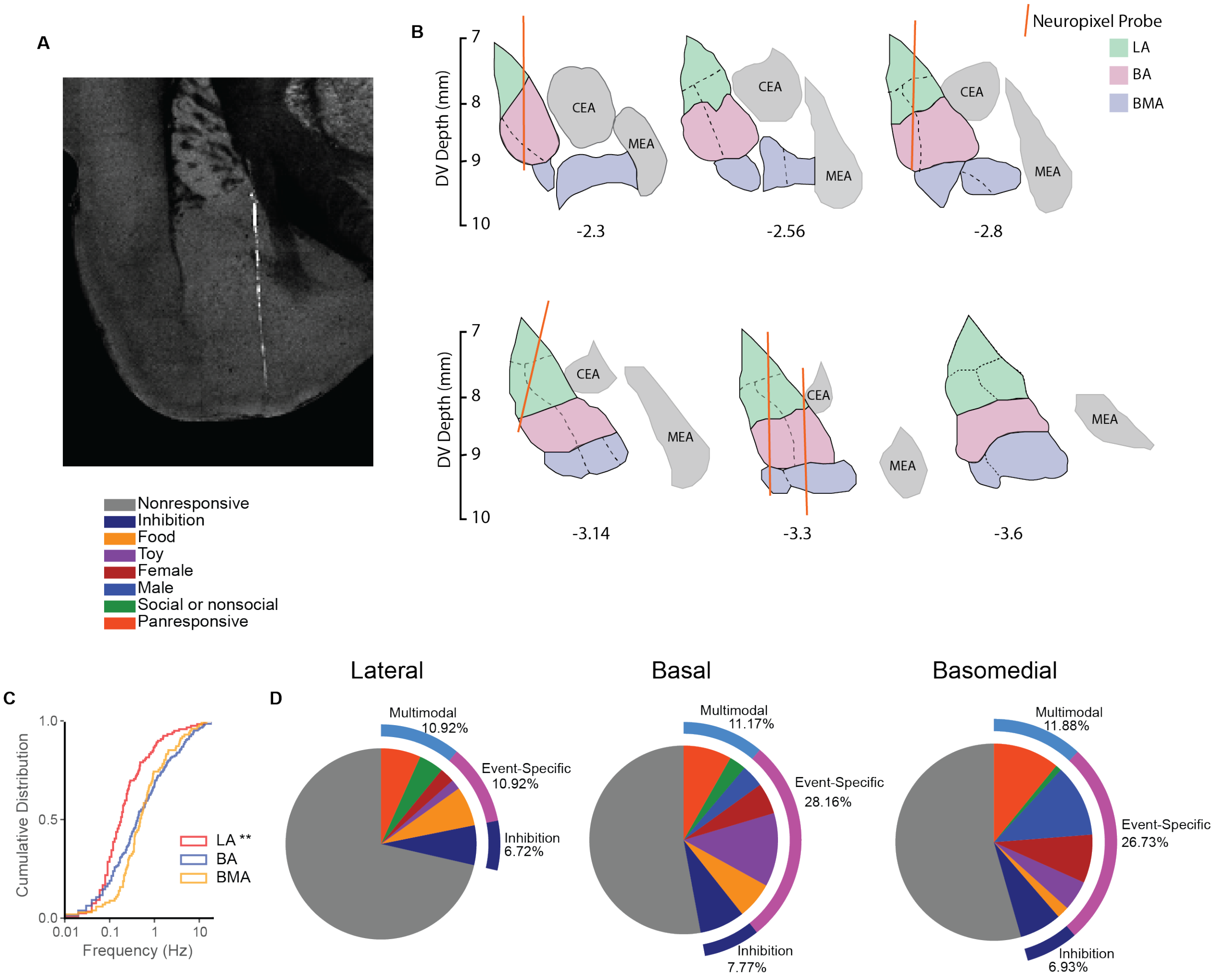


**Fig. S5. Histological Reconstruction of Neuropixel probe implants**

A) To visualize the electrode location, Neuropixel probes were dipped in Di-O prior to implantation. Following probe retrieval and perfusion, the brains were imaged using serial 2-photon tomography. Pictured is a representative image of a Neuropixels implant in the BLA. B) Neuropixel tracts were manually registered to a Paxinos rat brain atlas to determine which probe channels were located in specific BLA nuclei. C) LA has lower baseline firing rates than the BA and BMA. (**$p<0.01$ Kruskal Wallis test, LA significantly different from BA and BMA). This is consistent with the presence of fewer interneurons in that region (S3). D) LA (left) has fewer responsive neurons than BA(middle) and BMA (right)(see figure2) and neuronal selectivity to specific event-types varies across region with more non-social responses (i.e. event-specific food or toy) in the BA and more social responses in the BMA (i.e. event-specific male or female).

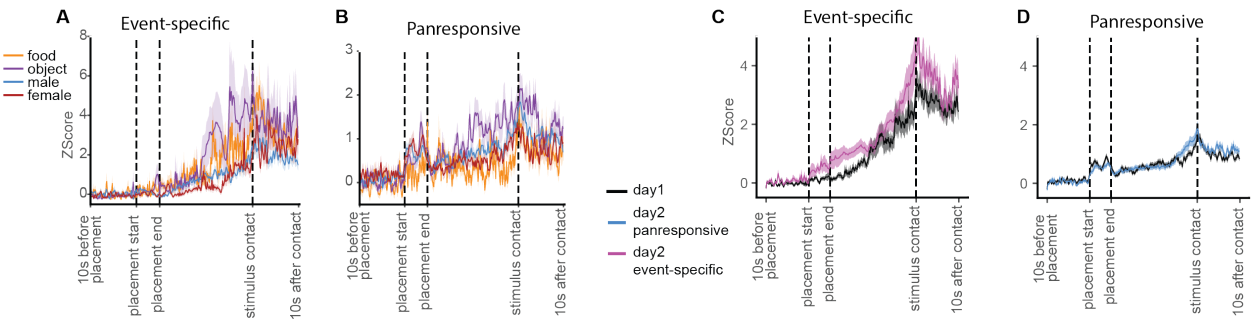


**Fig. S6. Similarities in firing onset between different events and across different recording days.**
A Event-specific and B) panresponsive neurons show similar time-courses irrespective of the stimulus identity, suggesting that the increases in firing are not due to the presence of a specific sensory cue but to occurrence of the event. Transient increases in firing in panresponsive neurons during stimulus presentation can be seen in all stimulus types. C and D) There were no differences in the timecourse in event-specific or panresponsive neurons across the two recording days.

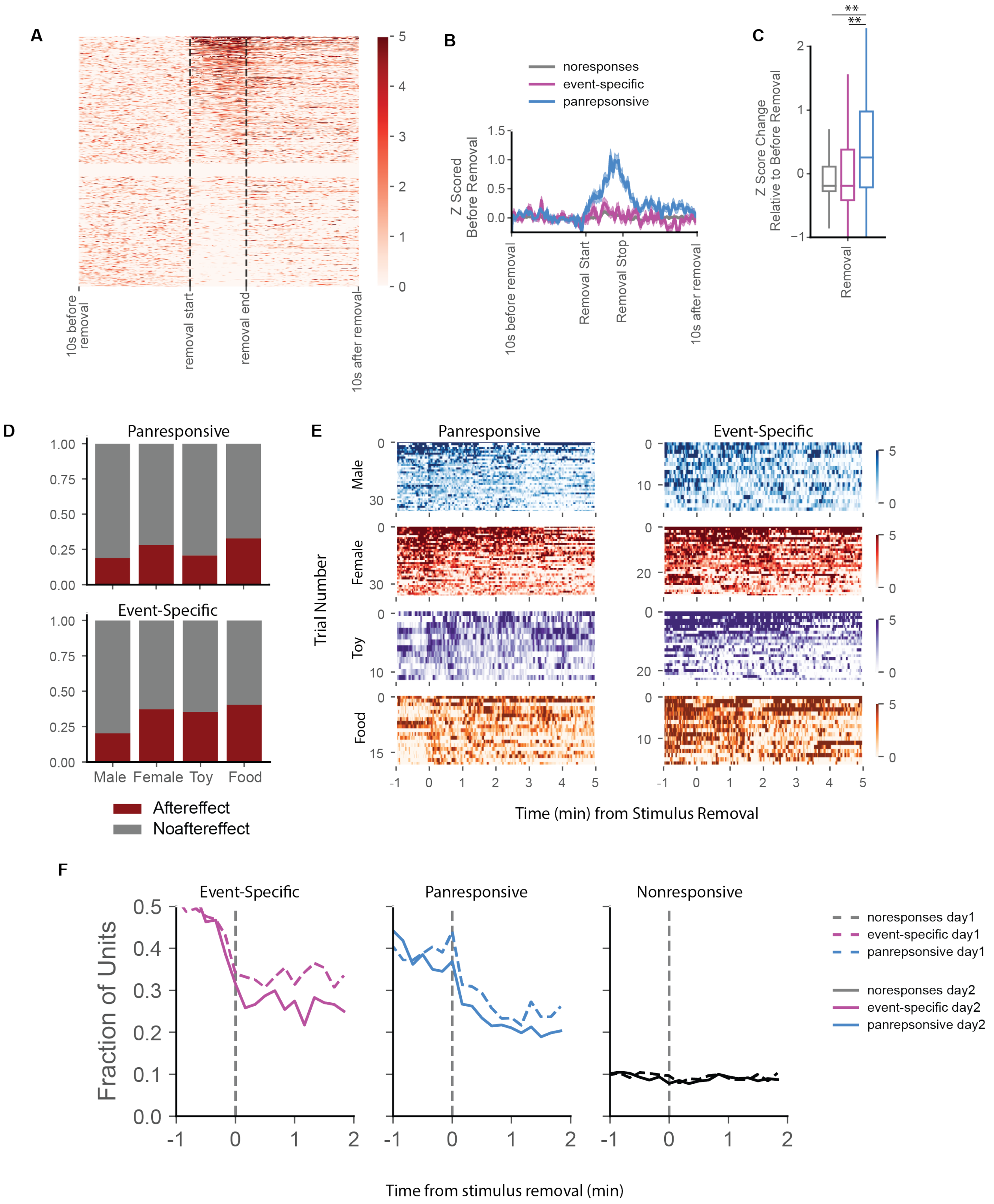


**Fig. S7. Neuronal firing persists after stimulus removal regardless of stimulus type**

A) Z-scored firing data from 10s before stimulus removal to 10s after removal. Data were timewarped between removal start and removal stop to enable comparison between different trials. B-C) Only panresponsive neurons show a transient increase in firing during the removal of the stimulus (**$p < 0.01$, ordinary one-way ANOVA for firing during removal period z-scored to previous 10s, N = 1239, 280, 389 for nonresponsive, event-specific and panresponsive trials respectively). D) The proportion of panresponsive and modality-specific neurons that show aftereffects is similar across different stimulus types. E) Z-scored images show after-effect producing neurons that continued to fire for up to 5 min following stimulus removal. F) The proportion of aftereffects found in event-specific and panresponsive neurons decreases slightly on the second day of recording.

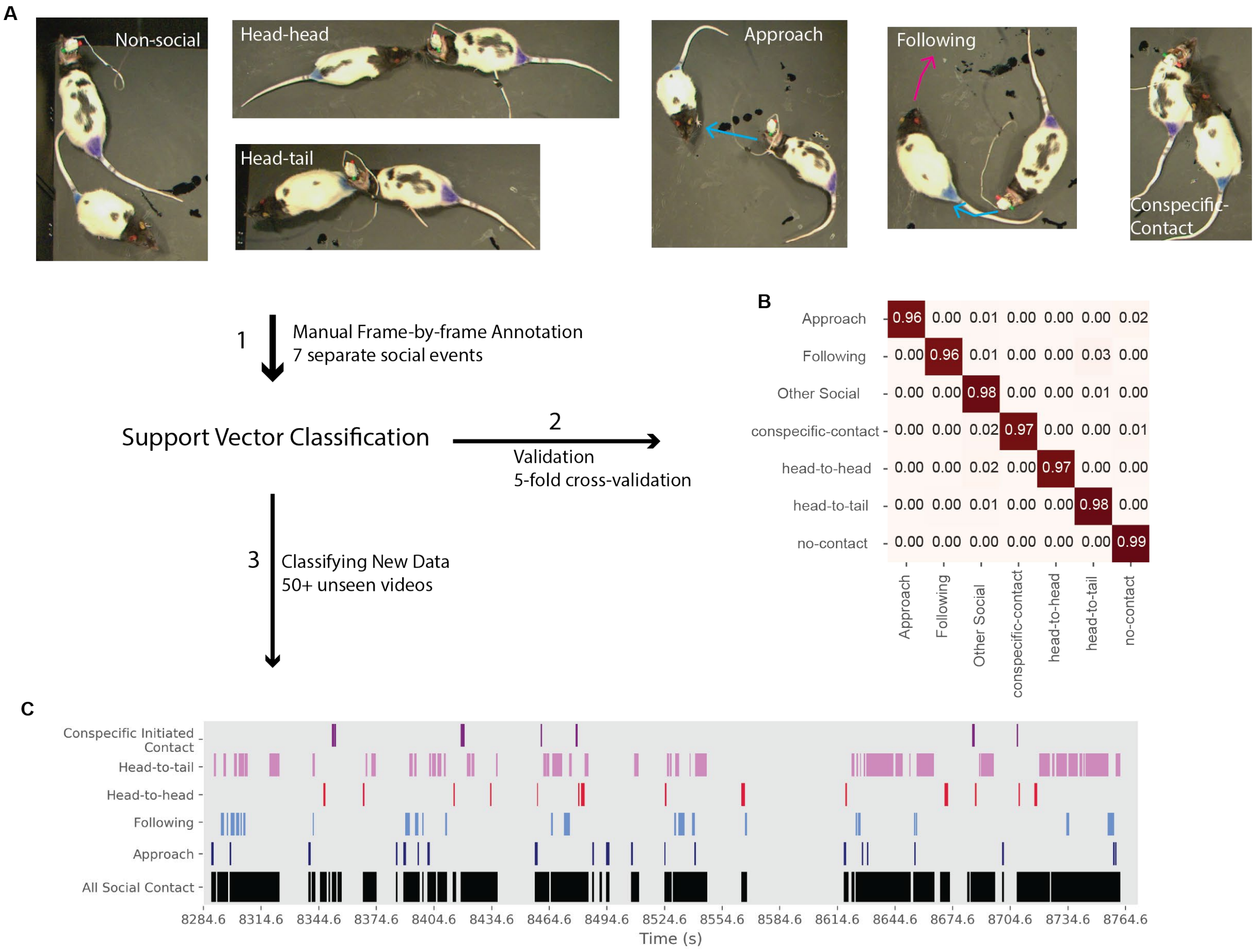


**Fig. S8. Automatic Classification of Social Behavior.**

A support vector classification was used to standardize the analysis of social behavior across all social interactions. A) 7 social interaction events (4 male-female events and 3 male-male events) were manually annotated frame-by-frame for one of 7 distinct behaviors (see methods for description of the behaviors). Pictured are non-social behavior and 5 distinct social behaviors. B) We trained the SVC classifier on the manually annotated behavior using the features extracted from the automated tracking. The SVC classifier was validated using 5-fold cross validation, which yielded balanced accuracies of 0.97. (see methods for a full-description of the analysis). C) We used the trained SVC classifier to automatically classify social behavior in previously unseen videos. Using this classification method, we identified two layers of behavioral activity from each social event. #1 – whether the animals were interacting (social) or not interacting (non-social) and #2 – the timing of 5 distinct social behaviors (sensory behavior – i.e. head-head or head-tail, movement behavior i.e. approach or following and passive i.e. conspecific contact).

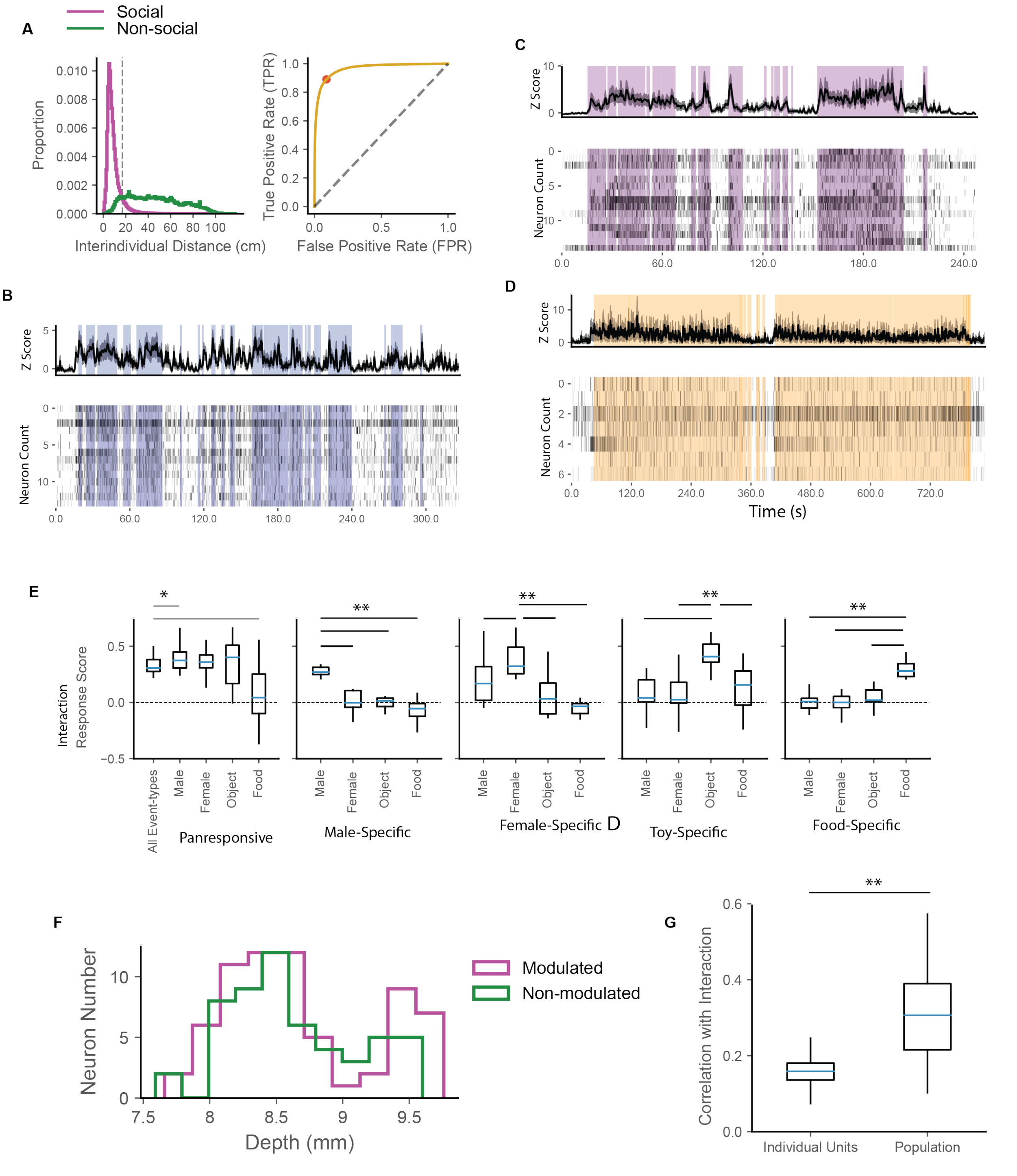


**Fig. S9. Properties of neurons modulated by interaction with stimulus.**

A) To automatically calculate stimulus-interaction in nonsocial events, we calculated the optimal interindividual distance threshold from all social trials using ROC analysis (see methods, threshold = 16.8cm). This threshold was also applied to non-social trials to identify behavioral interactions with the stimulus. B-D) Interaction-modulated neurons across different stimulus presentations illustrate the presence of single-cell and neuronal populations that reliably track engagement in both social and non-social stimuli B, males;  C, toy; D, food . E) Interaction-modulated event-specific neurons only show modulation by the preferred type of stimuli. For example, male-specific neurons do not show the same levels of modulation to female, toy or food stimuli. (** $p < 0.01$, repeated-measures one-way ANOVA with multiple comparisons, N = 8, 13, 15, 11 for male, female, toy and food engagement-modulated neurons). Panresponsive neurons show mixed selectivity (* $p < 0.05$, N = 15) F) Within responsive neurons, interaction-modulated neurons are not anatomically clustered compared to non-interaction modulated neurons. G) The population vector of interaction-modulated neurons is more highly correlated with behavior compared to the average individual units per event (** $p < 0.01$, Wilcoxon test N = 58 events).

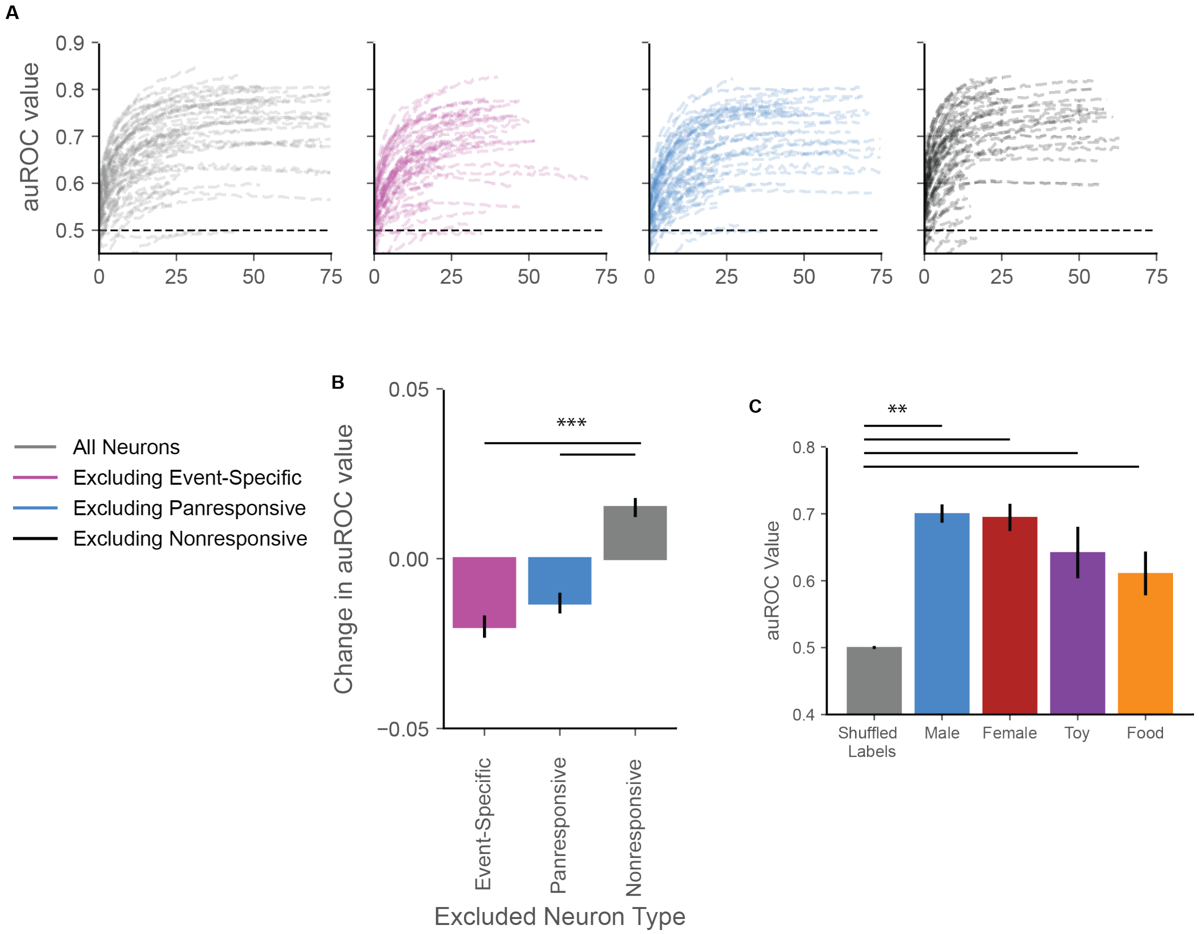


**Fig. S10. Decoding interaction with stimuli from neural activity**

A) Accurate decoding of behavioral interaction with stimuli is highly dependent on the number of neurons used (each line represents a single event). Removing specific neuron types (i.e. event-specific, panresponsive or nonresponsive) mildly impacts the performance of the decoder. This is quantified in (B, **p< 0.01, N = 60 events, one-way ANOVA with multiple comparisons). C) Behavioral interaction can be accurately decoded from event types with any of the 4 stimuli (**p<0.01, N = 60, 27, 17, 8, 8 for shuffled, male, female, toy, and food, Kruskal-Wallis test)

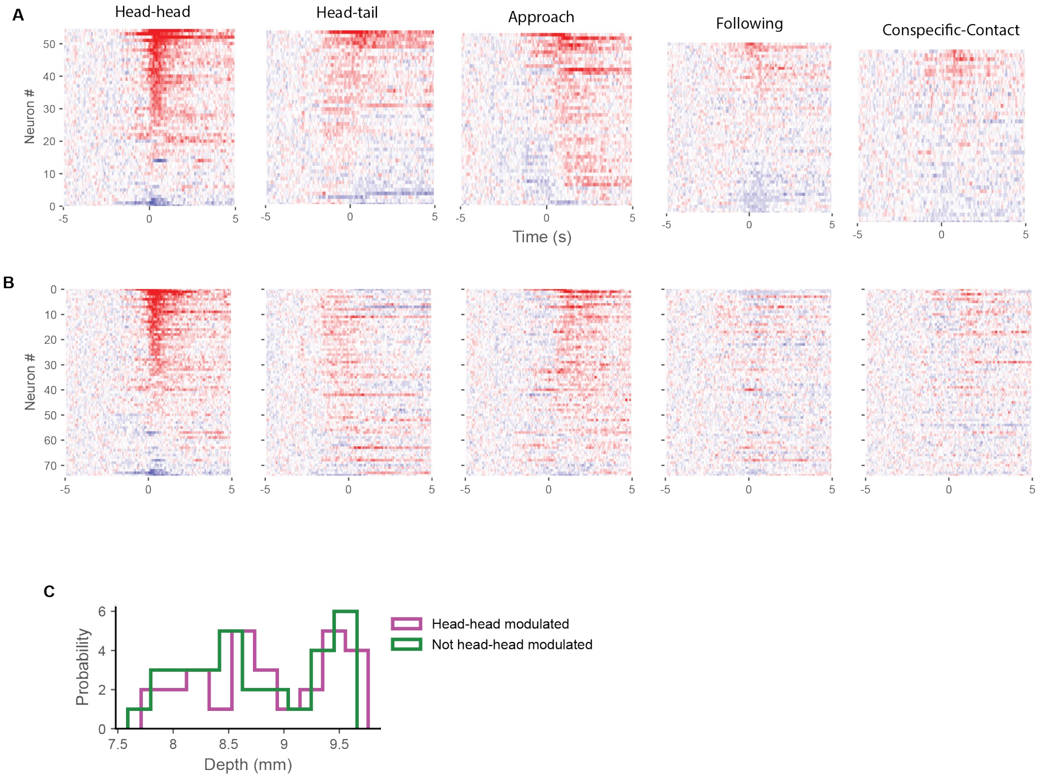


**Fig. S11. Properties of neurons modulated by specific social behaviors.**
A) A proportion of neurons show responses to specific social behaviors (each heatmap is sorted according to maximal response). More units respond to head-head, head-tail contact and approach than to following and conspecific-contact. B) Some units show selectivity to multiple social behaviors (all heatmaps sorted by the individual neuron's response to head-head contact). C) Head-head modulated neurons are not anatomically localized any more than non head-head modulated neurons.

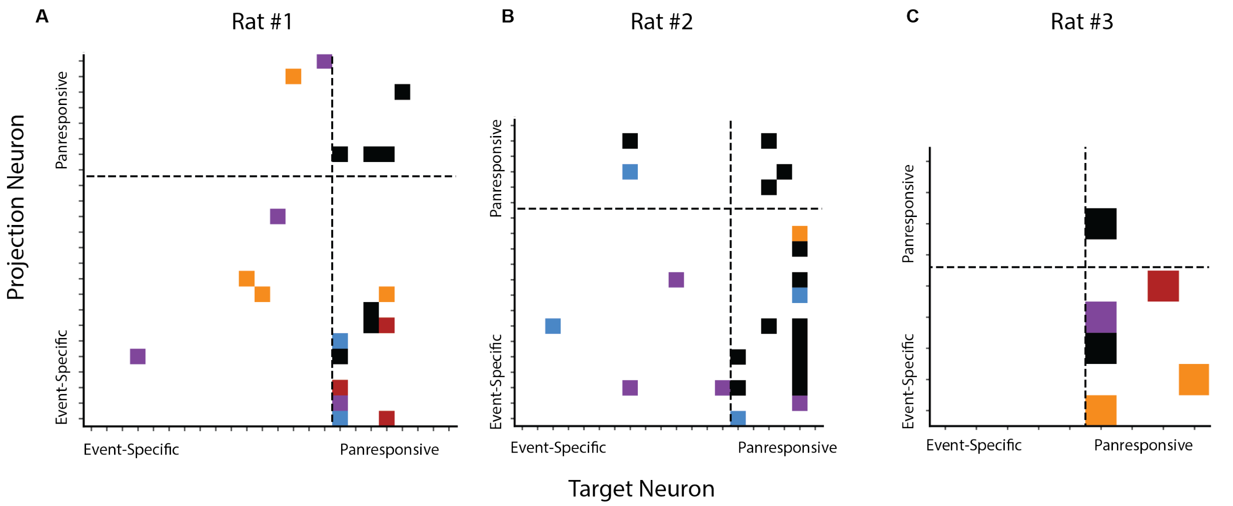


**Fig. S12.** Connectivity between event-specific and panresponsive neurons within individual rats. A-C) Cross-correlated pairs of neurons within 3 separate rats during the first presentation of each stimulus. Colors represent whether the neuronal pair fired exclusively during a single presentation (red, blue, purple, orange represents female, male, toy, food respectively) or if the neuronal pair was active across multiple stimulus presentations (black, 2+ stimulus presentations). During the stimulus presentation the flow of neuronal information typically goes from event-specific to panresponsive neurons.

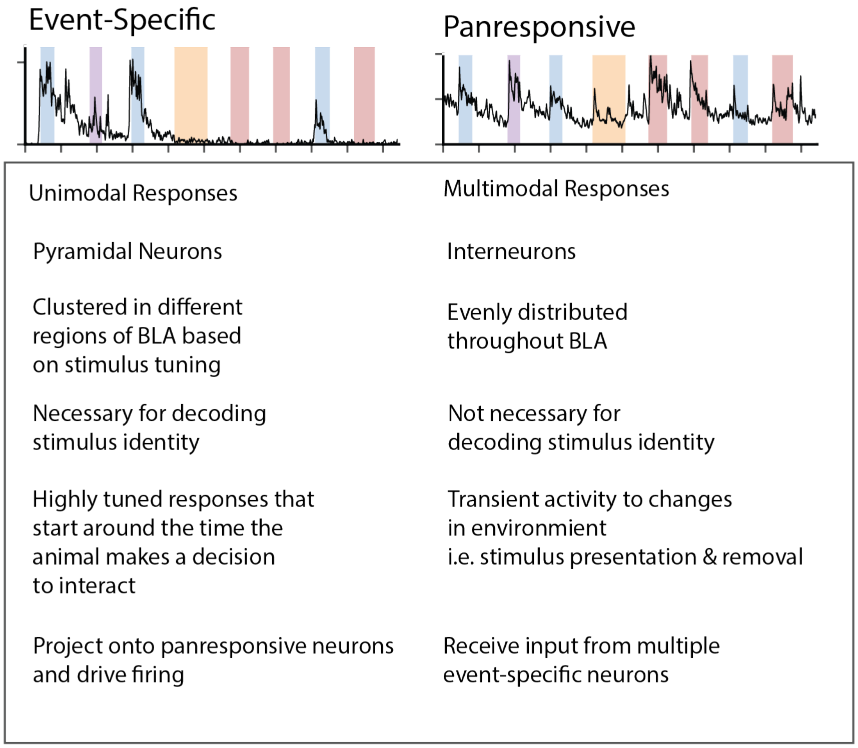


**Fig. S13. Summary of differences between event-specific neurons and panresponsive neurons.**

### Movie S1-S4.

An implanted rat interacts with another male conspecific (Movie S1), a female conspecific (Movie S2), a remote-controlled toy (Movie S3) or sweetened rice (Movie S4). Colored circles represent the coordinate locations of the tracked body parts (10 points per rat, 4 points for the toy and 1 for rice, with larger colored circles identifying experimenter colored bodyparts, see methods for more details).

**Movie S5**

Automated identification of social behavior during an interaction between the implanted male rat and a female rat. As described in FigS8 and methods, an SVC classifier was trained on manually annotated data and applied to previously unlabeled data like MovieS5.
